# Cryo-EM structure of the chain-elongating E3 ligase UBR5

**DOI:** 10.1101/2022.11.03.515015

**Authors:** Zuzana Hodáková, Irina Grishkovskaya, Hanna L. Brunner, Derek L. Bolhuis, Katarina Belačić, Alexander Schleiffer, Harald Kotisch, Nicholas G. Brown, David Haselbach

**Author notes:** These authors contributed equally (Zuzana Hodáková, Irina Grishkovskaya).

## Abstract

UBR5 is a nuclear E3 ligase that ubiquitinates a vast range of substrates for proteasomal degradation. This HECT E3 ligase has recently been identified as an important regulator of oncogenes, e.g., MYC, but little is known about its structure or mechanisms of substrate engagement and ubiquitination. Here, we present the cryo-EM structure of the human UBR5, revealing a building block of an antiparallel dimer which can further assemble into larger oligomers. The large helical scaffold of the dimer is decorated with numerous protein-interacting motifs for substrate engagement. Using cryo-EM processing tools, we observe the dynamic nature of the domain movements of UBR5, which allows the catalytic HECT domain to reach engaged substrates. We characterise the proteasomal nuclear import factor AKIRIN2 as an interacting protein and propose UBR5 as an efficient ubiquitin chain elongator. This preference for ubiquitinated substrates permits UBR5 to function in several different signalling pathways and cancers. Together, our data expand on the limited knowledge of the structure and function of HECT E3s.

## INTRODUCTION

Many vital signalling proteins, e.g., transcription factors and nuclear receptors, are key regulators of cell identity that are tightly controlled by multiple mechanisms, including their rapid degradation by the Ubiquitin-Proteasome System (UPS). Prior to proteasomal degradation, substrates often require a degradation mark in the form of ubiquitination which is achieved by a cascade of E1-E2-E3 enzymes. Many of these enzymes and the proteasome are enriched in the nucleus to swiftly respond to cellular signals^1^. One such enzyme is UBR5/EDD1/N-recognin-5, one of the largest human E3 ligases of the HECT class, initially identified as a tumour suppressor gene in drosophila^2^. However, several studies have since shown that UBR5 has a wide range of functions and substrates. Ubiquitination of substrates by this highly-conserved protein regulates events like mitotic progression by inactivation of the mitotic checkpoint complex^3,4^, transcription and DNA damage response^5–8^, or NF-κB immune signalling via the nuclear factor Akirin^9^. Its substrates extend far beyond those mentioned above (reviewed in ^10^). UBR5 also directly influences the cellular levels of multiple proto-oncogenes, such as p53^11^ or β-catenin^12^. Recently, we and two other groups described the great importance of UBR5 in cell growth via MYC degradation ^1,13,14^.

As many of these biological pathways are often implicated in cancer, it is not surprising that UBR5 was linked to numerous cancer types, including breast^15^, ovarian^16^, pancreatic^17^, colorectal cancers^18^ or mantle cell lymphoma^19^. In many cancer types, UBR5 is mutated, resulting in a loss of its tumour-suppressing activities via deregulation of DNA repair pathways or directly influencing the levels of proto-oncogenes^13^. In contrast, UBR5 also has additional oncogene-like functions, for example, by enhancing tumour growth through inhibition of the cytotoxic T-cell response or modulating tumour microenvironments through immunosuppression^20,21^.

How UBR5 regulates such a vast range of substrates and signalling pathways is poorly understood. The current structural information is limited to a few crystal structures of isolated domains^22,23^, leaving the majority of UBR5 structurally uncharacterised. At the UBR5 C-terminus lies the highly conserved HECT catalytic domain, which mediates the transfer of ubiquitin from the E2 enzyme onto itself before attachment to the target substrate. The HECT domain is a conserved feature of all 28 E3 ligases of the HECT class^24^. While studies focused on isolated domains of HECT E3 ligases shed some light on their function^25,26^, solving the full-length structure of the first HECT-type E3 ligase, HUWE1, revealed that the complex architecture plays a key role in substrate engagement in relation to ubiquitin transfer^27,28^. Yet, there is considerable heterogeneity between HECT enzymes, and more structural and biochemical information is needed to understand the polyubiquitination mechanism of UBR5 and HECT E3s in general.

To determine the potential interplay between the catalytic HECT domain and protein-interacting domains, we solved the cryo-EM structure of full-length human UBR5. We show that the protein-protein interaction domains are inserted in the large armadillo-type helical scaffold. Excitingly, UBR5 assembles into homodimers, tetramers, and even larger oligomers. This striking feature allows the positioning of substrate binding sites in close proximity to the catalytic HECT domain in cis or trans, expanding its substrate-recruiting capacities. In addition, we uncovered multiple protein binding domains and identified an interacting partner, AKIRIN2. Using AKIRIN2 as a substrate, we find that UBR5 exhibits a preference to extend ubiquitin chains from a pre-ubiquitinated substrate, suggesting UBR5 functions primarily as a chain-elongating E3 ligase. The structural and biochemical information presented begins to unravel the versatility of UBR5 for its substrate choice and function.

## RESULTS

### UBR5 dimer is the functional unit

UBR5 is a highly conserved protein and is almost completely restricted to metazoans. Human UBR5 is 2799 residues long, and this length is an average across all species (Suppl.Table 1). Because of its high conservation, we focused on obtaining the structure of the full-length human UBR5 protein. We expressed UBR5 in a stable HEK293T cell line and purified it using affinity and size exclusion chromatography (SEC). SEC elution profile revealed that UBR5 elutes as two peaks. Using mass photometry (MP), a method for the determination of heterogeneous species and oligomers found in an experimental sample^29^, we found that UBR5 assembled into oligomers in both SEC elution peaks (Figure 1B, Suppl.Figure 1A,1B). Interestingly, the oligomers observed were formed in a repeating order of two, suggesting that a dimer is the building block and functional unit of all oligomers.

**Figure 1.**
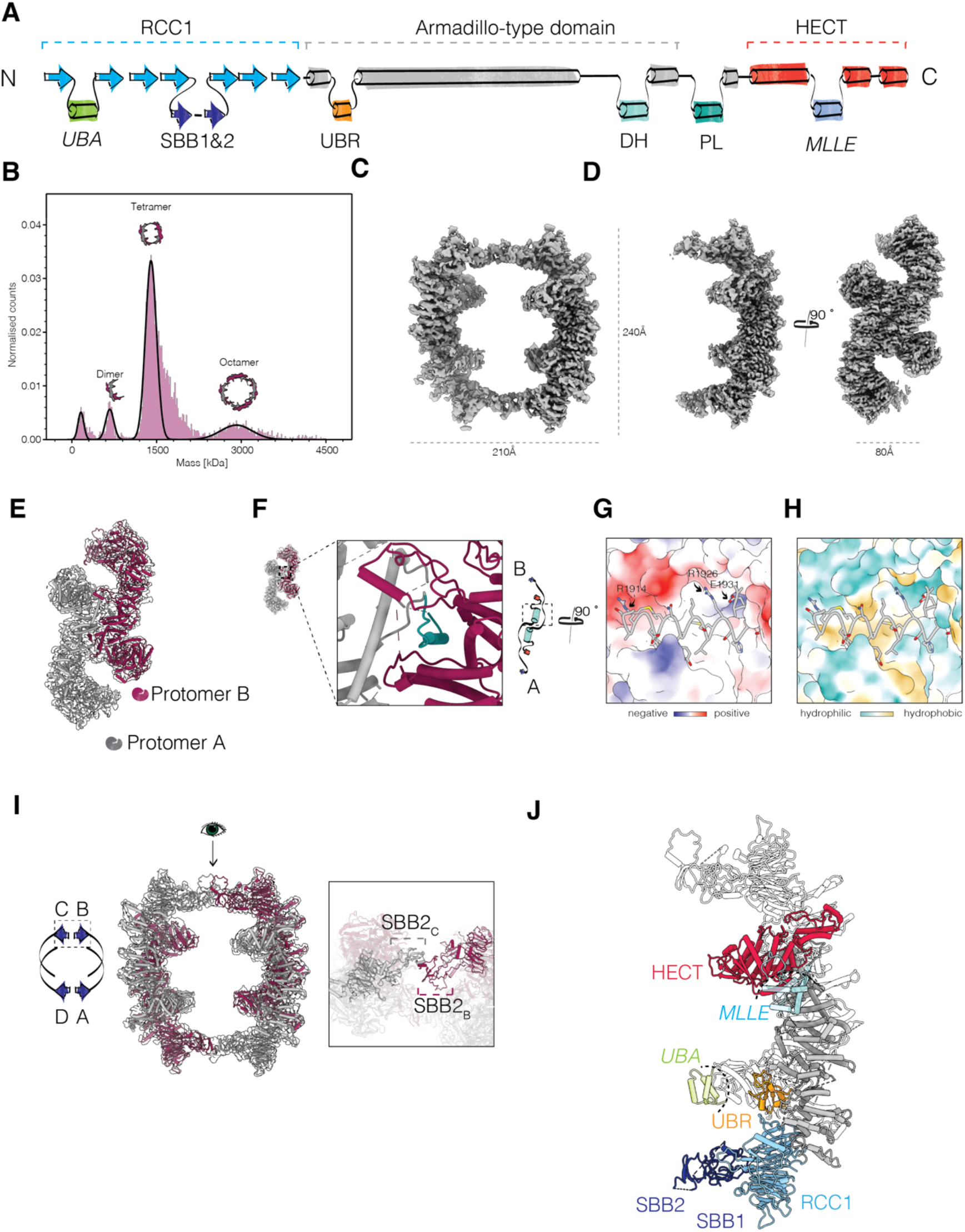
Cryo-EM structures of the dimeric and tetrameric UBR5 assemblies. A) Schematic representation of domain organisation of the human UBR5 protein. UBA – Ubiquitin associated domain, RCC1-Regulator of chromosome condensation 1, SBB – small β barrel, DH - dimerisation helix, PL – plug loop, MLLE – Mademoiselle. Mass distribution and oligomeric states adopted by UBR5, measured using mass photometry. C) Single particle cryo-EM reconstruction of the tetrameric UBR5. D) High-resolution cryo-EM reconstruction of the dimeric UBR5. E) SBB domain interactions between two UBR5 dimers. F) Molecular details of the dimerisation interface of two UBR5 protomers in one dimer unit. G) Atomic model of the UBR5 homodimer, colour-coded in relation to (A). For clarity, only one protomer is highlighted. Positions of unresolved domains were estimated based on AlphaFold2 predictions. The unresolved domains are depicted in opaque colours, annotated in italics.

Our attempts to separate species found in individual peaks were unsuccessful. The second SEC elution peak was less heterogeneous, consisting predominantly of tetramers, and was thus chosen for structural analysis using single-particle cryo-electron microscopy (cryo-EM). 2D classification and subsequent processing revealed that UBR5 formed a large ring-shaped assembly with overall dimensions of 215×215×90Å and a large central cavity (Figure 1C, Suppl.Figure 2B). We used AlphaFold2 to predict the structure of UBR5 and docked it into our cryo-EM map, which occupied approximately a quarter of the density, consistent with our observations that our input UBR5 sample formed tetramers. While Alphafold2 did not successfully predict UBR5 tetramers or dimers, docking in multiple copies of the predicted structure confirmed that the ring represents a tetrameric assembly of UBR5. Our initial dataset suffered from insufficient particle orientations to confidently build an atomic model. We, therefore, collected and merged several untilted and tilted UBR5 datasets, which corrected for map anisotropy (Suppl.Figure 2). Additionally, refinements of the initial tetrameric map always converged with one half of the map resolving significantly better than the other, which we interpreted as flexible motion between the two halves. To cope with this conformational flexibility, we performed local refinements using a half-map mask. This approach allowed us to obtain a high-resolution cryo-EM map, which clearly showed that each half of the ring is formed by two UBR5 protomers arranged in an antiparallel fashion (Figure 1D,1E, Table1).

**Table1.**
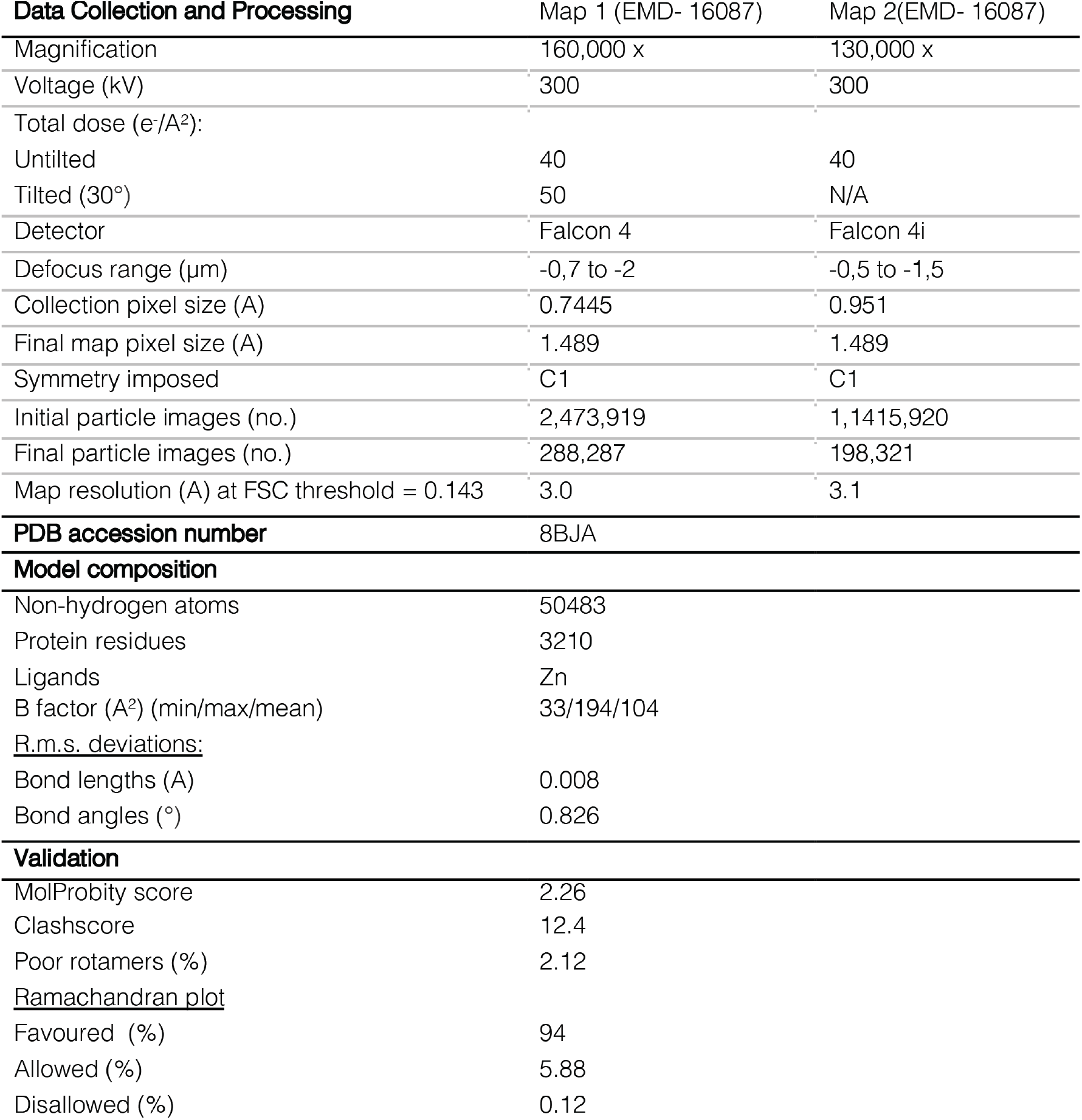
Cryo-EM data collection, refinement, and validation statistics.

| **Data Collection and Processing** | Map 1 (EMD- 16087) | Map 2(EMD- 16087) |
| --- | --- | --- |
| Magnification | 160,000 x | 130,000 x |
| Voltage (kV) | 300 | 300 |
| Total dose (e-/Å<sup>2</sup>): |  |  |
| Untilted | 40 | 40 |
| Tilted (30°) | 50 | N/A |
| Detector | Falcon 4 | Falcon 4i |
| Defocus range (µm) | -0,7 to -2 | -0,5 to -1,5 |
| Collection pixel size (Å) | 0.7445 | 0.951 |
| Final map pixel size (Å) | 1.489 | 1.489 |
| Symmetry imposed | C1 | C1 |
| Initial particle images (no.) | 2,473,919 | 1,1415,920 |
| Final particle images (no.) | 288,287 | 198,321 |
| Map resolution (Å) at FSC threshold = 0.143 | 3.0 | 3.1 |
| **PDB accession number** | 8BJA |  |
| **Model composition** |  |  |
| Non-hydrogen atoms | 50483 |  |
| Protein residues | 3210 |  |
| Ligands | Zn |  |
| B factor (Å<sup>2</sup>) (min/max/mean) | 33/194/104 |  |
| <u>R.m.s. deviations:</u> |  |  |
| Bond lengths (Å) | 0.008 |  |
| Bond angles (°) | 0.826 |  |
| **Validation** |  |  |
| MolProbity score | 2.26 |  |
| Clashscore | 12.4 |  |
| Poor rotamers (%) | 2.12 |  |
| <u>Ramachandran plot</u> |  |  |
| Favoured (%) | 94 |  |
| Allowed (%) | 5.88 |  |
| Disallowed (%) | 0.12 |  |

**Figure 2.**
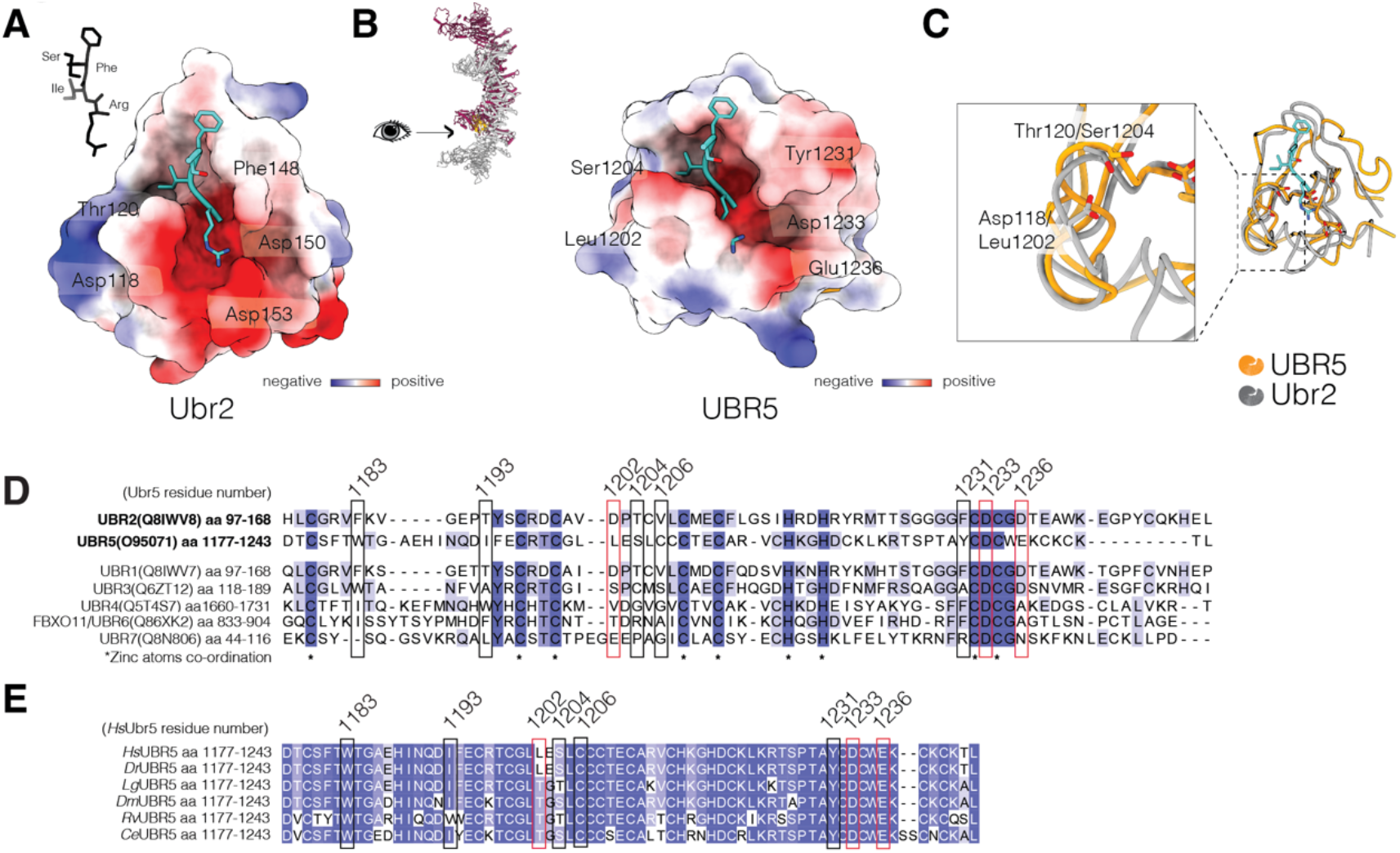
The structure and sequence conservation of the UBR box. A) Surface representation of the electrostatic potential of the UBR2 UBR box (PDB:3NY3) bound by an Arg/N-degron mimic peptide, “RIFS”. Key residues forming the interaction with the arginine and isoleucine of the peptide are highlighted. B) Surface representation of the electrostatic potential of the UBR5 UBR box. The RIFS peptide was positioned into the UBR5 UBR box by aligning the UBR2 UBR box crystal structure to the UBR5 model (see C). UBR5 residues corresponding to the residues mediating the N-degron binding in UBR2 are highlighted. C) An overlay of the UBR2 (PDB:3NY3) and UBR5 UBR boxes. UBR2 UBR box is bound by the peptide RIFS. Highlighted in the dashed box is the distinct fold of the two boxes. Highlighted residues represent amino acids which mediate binding to RIFS in UBR2 but do not have conserved properties in UBR5. D) Sequence comparison of UBR boxes across 7 human UBR E2 ligases. Residues mediating the contacts with the N-degron are highlighted in boxes (in red – contacts with the terminal arginine residue, in black – contacts with the penultimate hydrophobic residue) numbered according to the UBR5 sequence. E) Sequence conservation of the UBR5 UBR box across species. Residues predicted to mediate contacts with the N-degrons are highlighted in boxes, colour-coded as in D.

### UBR5 homodimers are decorated with substrate-recognition domains

Using the AlphaFold-predicted UBR5 model as a template for model building, we were able to confidently model the UBR5 dimer, characterise the dimerisation interface and assign densities of several protein binding domains of UBR5 (Figure 1A, 1G). The central feature of the structure is a large alpha-helical scaffold that has previously not been predicted for UBR5 ^24^. The scaffold, assembled into an armadillo-type fold domain, spans approximately 100Å between the N and C termini of UBR5.

The most striking feature of our cryo-EM density was the elegant assembly of UBR5 into an antiparallel homodimer. The interface is mainly mediated by a single helix extended from one protomer into the other (Figure 1F-1H). This heavily conserved interaction interface is formed by a single helix of one protomer inserting itself into a hydrophobic pocket of the opposite protomer. The hydrophobic cavity of the pocket is flanked by several charged residues on each side (R1914, R1926, E1931), forming salt bridges with residues of the opposite protomer (D1381, R1434 and D1505, respectively). Our findings that the dimer as a functional unit remains intact even in high salt further corroborates this finding (Suppl.Figure 1C).

The helical scaffold and the N terminus of UBR5 are composed of numerous protein-interacting motifs (Figure 1A, 1J). The antiparallel assembly of the UBR5 dimer places these motifs into relatively close proximities to the catalytic HECT domains on the C termini of the protomers. The N-terminus revealed a complex spatial arrangement of three protein-interacting motifs. The absolute N-terminus consists of a 7-bladed β -propeller domain that has previously been annotated as an RCC1-repeat domain^24^. Fitting in the crystal structure of the RCC1 protein revealed a near-identical conformation^30^. A Similar was observed when fitting in the RCC1 domain of HERC2, a HECT-type E3 ligase. The RCC1 domain is discontinuous; it contains three accessory domain insertions. After the first blade of the β-propeller, a long segment of disordered sequence spanning almost 300 amino acids exits the RCC1 domain and inserts back to continue into the second blade(Figure 1A, 1J). Within this loop is nested the UBA domain, but because of its flexible tether, we cannot predict the position of the UBA domain. It is likely that these loops allow the UBA domain to be dynamic and possibly utilise proximity-based interactions.

The second two insertions are inserted just after the 4^th^ blade of the RCC1 domain. Due to resolution limitations in our cryo-EM reconstructions, we can assign the relative positions of these two domains but not assign secondary structures. These domains are predicted to be small domains with a five β-strand assembly. Based on their appearance, we classify them as small β-barrels (SBBs). These domains are often directly involved in RNA binding or serve as assembly platforms for RNA-associated protein complexes. Additionally, numerous SBB domains were found to facilitate oligomerisation ^31^. We performed extended 3D classification of the tetramer, which allowed us to confidently dock two models of UBR5 dimers. This revealed that the tetrameric assembly of UBR5 is formed by SBB2 domains of two opposite dimers(Figure 1I). Additional 3D variability analysis (3DVA) ^32^ revealed a dynamic movement between two opposing dimers(Movie 1). The flexibility appeared to stem from the SBB2 domain contacts. The interactions are most likely transient, allowing UBR5 to oligomerise into higher multimers. Importantly, such movements could also bring the catalytic and protein-binding motifs closer together within the lumen of the ring.

Another insertion of the helical scaffold is the UBR box, a key feature giving UBR5 its name and placement into the family of UBR E3 ligases/N-recognins^33^. The UBR box is a conserved substrate recognition domain of all members of the UBR family^34^. The UBR box of UBR5 resembles the structures of several other published UBR boxes ^35,36^. The box is stabilised by three zinc ions mediated by two histidine and seven cysteine residues, which are absolutely conserved across all UBR boxes(Figure 2C). UBR domains function as binding domains for N degrons: substrates with a destabilising N terminal amino acid. UBR5 was, similarly to UBR1,2 and 4, reported to bind type-I Arg/N-degrons, which contain positively charged terminal residue and a hydrophobic penultimate residue. These bind to a negatively charged pocket and a secondary hydrophobic pocket in the UBR box ^36–38^(Figure 2A). Overlays of Ubr2 and UBR5 UBR boxes and docking in the degron-mimicking peptide “RIFS” into UBR5 UBR box clearly shows a change in the overall domain fold, despite several conserved or similar residues forming the degron-binding pockets (Figure 2B-2D). The negatively charged pocket in UBR5 is less prominent than in Ubr1 or Ubr2, formed by two acidic residues (D1233 and E1236) instead of three. The third is swapped for a hydrophobic leucine (L1202) across all species (Figure 2E). This possibly extends the hydrophobic pocket binding the penultimate degron residue. Overall, this secondary pocket is well-preserved in UBR5. Together the changes in the surface-exposed residues and the overall fold of the UBR box could imply differences in N-end rule substrate selection between UBR5 and other types of Arg/N-degron-binding E3 ligases.

UBR5 contains one additional protein-interacting motif, the MLLE domain. Whilst we cannot resolve this domain due to its flexible nature, we can trace the region in the N-lobe, which extends into the linkers between the HECT and MLLE domains. Contrary to the previous annotations^24^, this motif does not precede the HECT domain but is rather an insertion into the N-lobe, being an uncommon characteristic for HECT domains. The linkers connecting MLLE and HECT domains span approximately 40 residues on either side of MLLE and could aid in MLLE-bound substrate transfer to the HECT domain. Interestingly, one of the linkers includes the sequence previously found to be a PAM2-like sequence, which can bind the MLLE domain directly as an isolated peptide^22^. Whether the MLLE domain interacts with the HECT domain in the context of the full-length protein remains to be investigated.

### UBR5 is an efficient ubiquitin chain extender

We next focused on characterising the ubiquitination activity of UBR5. Previous reports suggest that the homolog of UBR5 in *D. melanogaster*, Hyd, ubiquitinates Akirin in the NF-κB signalling^9^. In humans, there are two orthologs of Akirin, AKIRIN1 and AKIRIN2. We have previously shown that AKIRIN2 is essential for the turnover of the oncogene MYC as well as virtually all nuclear proteins via facilitating the nuclear import of the proteasome^1^. The screen which identified AKIRIN2 as a regulator of MYC also identified UBR5 as one of the E3 ligases governing its turnover whilst not being essential.

We, therefore, hypothesised that UBR5 might interact and ubiquitinate AKIRIN2 as seen in other systems. We performed activity assays using a GFP-labelled AKIRIN2 and two control substrates, Securin and GFP (Figure 3A, Suppl.Figure 3B, 3D). We chose Securin as it has the identical length of AKRIN2 and is similarly unstructured. However, we did not observe any ubiquitination of either substrate but have observed the formation of free ubiquitin chains. In parallel, we tested whether AKIRIN2 binds UBR5. We tested their interaction using sucrose gradients and observed a co-migration pattern, showing that they interact *in vitro*. Importantly, Securin nor GFP co-migrated with UBR5 (Figure 3B, Suppl.Figure 3C, 3D).

**Figure 3.**
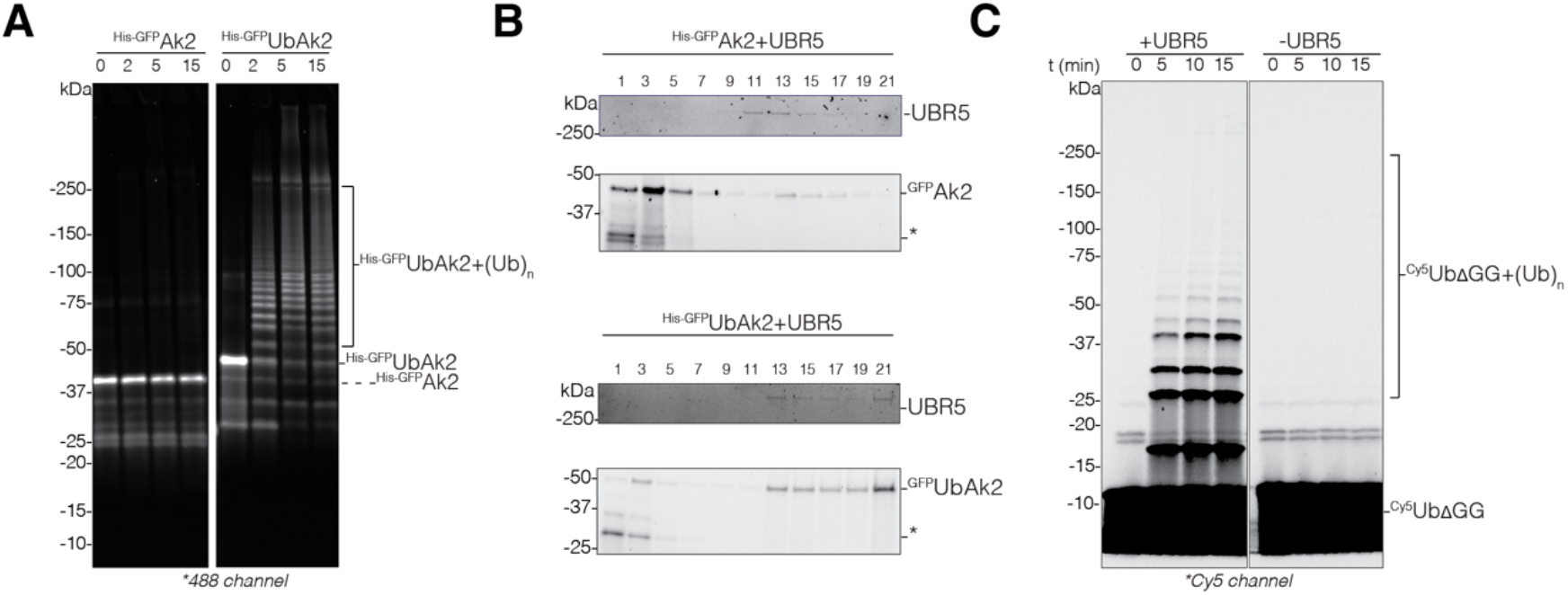
Substrate selection of UBR5. A) Ubiquitination assays of an unmodified and monoubiquitin-like substrate. Ubiquitinated products were visualised by imaging the GFP used to tag substrates. B) Sucrose gradient sedimentation profiles of UBR5 mixed with two substrates. UBR5 was visualised using stain-free imaging. Substrates were visualised by imaging their GFP tag. C) Ubiquitination assays of UbΔ^GG^ in the presence and absence of UBR5. Ubiquitinated products were visualised by imaging the Cy5 fluorophore used for labelling UbΔ^GG^.

Because AKIRIN2 bound UBR5, we considered that the inability of UBR5 to ubiquitinate AKIRIN2 might stem from poor chain-initiating properties of the enzyme. Instead, we anticipated UBR5 might be effective as a chain elongator with a pre-requisite of substrates with a ubiquitin modification. It was previously suggested that UBR5 co-operates with other E3s to extend chains, for example, with ITCH, UBR4 and HUWE1, for TXNIP ubiquitination and proteotoxic stress response ^39,40^. We, therefore, expressed AKIRIN2 N-terminally fused to ubiquitin (UbAKIRIN2) and tested both its ubiquitination by and binding to UBR5. Additionally, we confirmed our selection of the E2-conjugating enzyme, UBCH5B, using the E2 screening kit to exclude the possibility of the wrong selection of enzyme partners(Suppl.Figure 3A). Using UbAKIRIN2, We observed rapid growth of ubiquitin chains on the substrate. Interestingly, we observed a similar profile when we fused ubiquitin to Securin (UbSecurin). We, therefore, repeated sucrose gradient experiments using the two monoubiquitinated substrate mimics. We observed no binding of UbSecurin (Suppl. Figure 3C). Contrary to this observation, the interaction between UBR5 and UbAKIRIN2 was enhanced by the presence of ubiquitin in the construct. These findings suggest an avidity mechanism of engagement where the substrate first binds to a specific protein-interacting motif on UBR5 and is subsequently engaged by, presumably, the UBA domain, which binds ubiquitin. However, certain interactions could be very transient, especially in the case of substrate ubiquitination. Because we observe ubiquitination of UbSecurin, it is possible that there are very transient, possibly proximity-mediated, interactions with the enzyme. In the case of AKIRIN2, the interaction may serve other functions where a more stable binding is required.

We further confirmed the chain-elongating properties of UBR5 by using a Cy5-labelled ubiquitin with a deletion of the two C-terminal glycine residues (Ub^ΔGG^). This ubiquitin, serving as a ubiquitin acceptor, was used instead of a substrate in activity assays. Ub^ΔGG^ was efficiently ubiquitinated by UBR5, where the presence of high molecular weight species suggests that it was decorated by polyubiquitin chains, as the molecular weight cannot be achieved only by monoubiquitination at distinct lysines of Ub^ΔGG^. The ubiquitination properties could be attributed to UBR5, as in its absence, no ubiquitination was observable (Figure 3C). We confirmed the linkage type using mass spectrometry and found that UBR5 formed almost exclusively K48-type ubiquitin linkages *in vitro* (Suppl.Table2).

### Interfaces mediating the HECT domain and UBR5 activity

The HECT domains of all described HECT E3 ligases lie in the absolute C-terminus of the protein. Structurally, these domains have a conserved architecture composed of N- and C-lobes. The N-lobe facilitates E2-ubiquitin binding whilst the C-lobe mediates the transfer of ubiquitin first onto itself and then onto a target substrate via its catalytic cysteine residue ^24^.

For the HECT domain to ubiquitinate substrates bound to the binding modules of UBR5, a certain degree of flexibility is required. In addition to the movement of the whole helical body of the dimeric and tetrameric assembly, we observed a movement of the HECT domain with respect to the rest of the protein (Movie 2). We identified two states: in state 1, the HECT domain is positioned facing downwards, whereas in state 2, the domain is rotated, and the C-lobe is shifted by 30**°** upwards (Figure 4A). State 2 is less abundant in our datasets, with only about a fourth of particles adopting this state. The movement of the HECT domain is, however, visible amongst the 2D class averages (Figure 4B). In HUWE1, the flexibility of the HECT domain is attributed to the hinge-like set of three helices emerging from the helical scaffold to the base of the N lobe (Figure 4C). Mutations in the hinge were found to significantly impact ubiquitination activity of HUWE1, suggesting the HECT dynamics is crucial for its enzymatic activity^27,28^. A three-helix bundle hinge domain in UBR5 with two helices positioned identically to HUWE1 divides the HECT domain and the armadillo repeats of UBR5, forming the only continuous contact between the enzyme body and the catalytic domain. It is, therefore, probable that the flexibility of the HECT domain is a requirement for UBR5’s ubiquitination activity and likely numerous other HECT-type E3 ligases.

**Figure 4.**
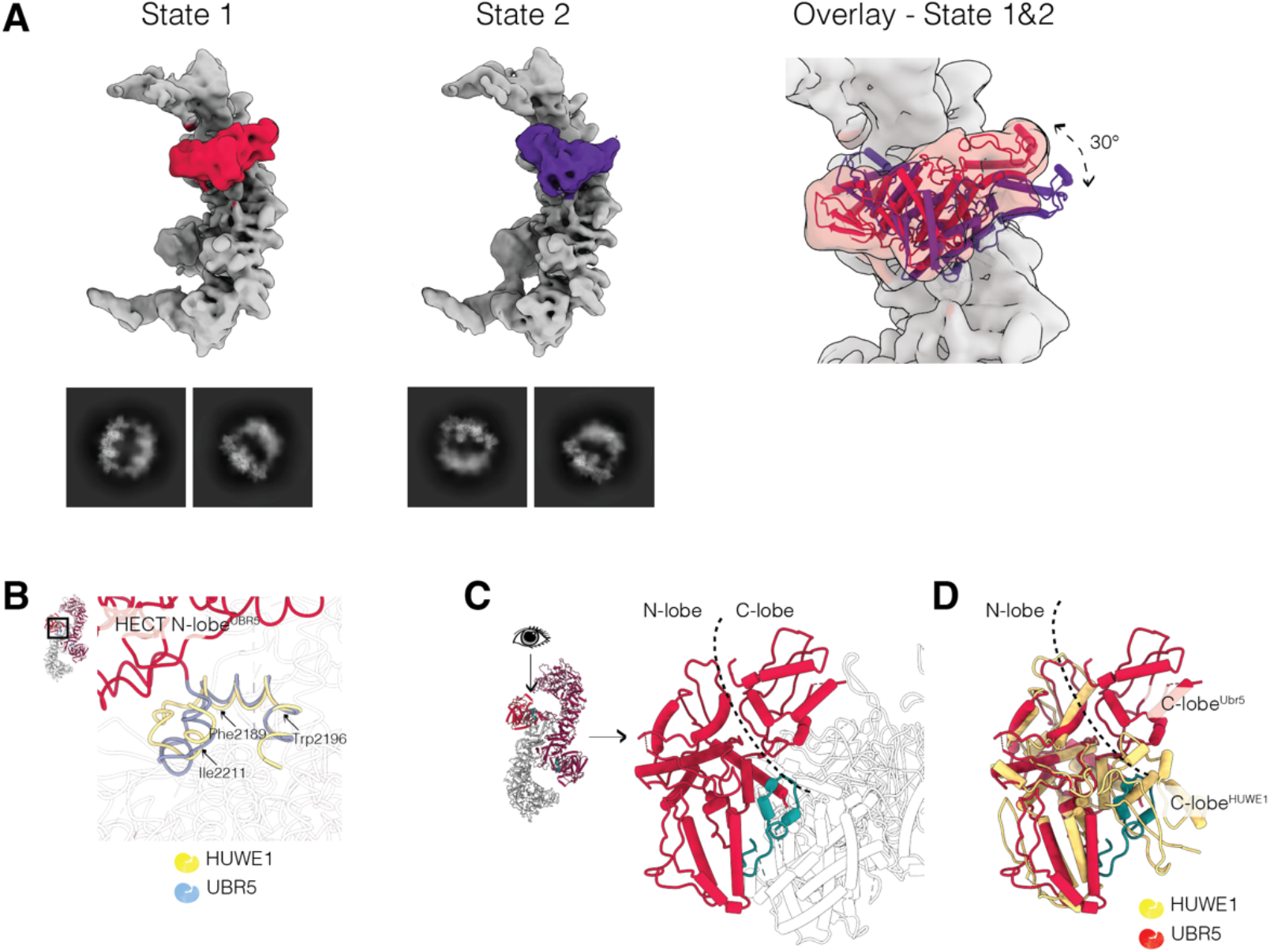
HECT domain orientations. A) Conformational states of the HECT domain identified using 3DVA with an overlay of the two states with the UBR5 HECT domain fitted into state1. A 30degree rotation is observable between the two states. B) An overlay of the hinge loop of HUWE1 (in gold) and UBR5 (in blue). Residues of UBR5 predicted to govern the same function as in HUWE1 are highlighted. C) Top view of the HECT domain (in red) and the plug loop in teal. D) Overlay of the HECT domains of human HUWE1 (PDB:7MWD, in gold) and UBR5 (in red). The HECT C-lobe of HUWE1 overlaps with the position of the plug loop (in teal) found in UBR5.

We additionally identified another region, termed the plug loop, a stretch of conserved 25 amino acids extending from the armadillo repeats between the lobes of the HECT domain before looping back to the repeats (Figure 4D). This loop does not precede the hinge or HECT domains in a linear sequence but rather is separated by almost 200 unstructured residues. Aligning UBR5 with the full-length HUWE1 structure reveals that the observed positions of the two HECT lobes are significantly different (Figure 4E). There is no loop occluding the space between the HECT lobes of HUWE1, and therefore they are in a compact, T state. In UBR5, the HECT lobes are spatially separated by the plug loop. We speculate that conformational rearrangements in UBR5 are necessary to disconnect the plug and allow for C-lobe rotation for ubiquitin transfer.

### A model for substrate engagement and ubiquitin transfer

The observed states of the HECT domain, together with the possible inhibitory effect of the plug loop, raise the question of what steps are necessary for ubiquitin transfer. Whilst it is clear that the plug loop must be removed from its stationary position to allow for C lobe rotation, both HECT domain states observed in our datasets appear to be in an open, L conformation. These states could potentially represent pre-transfer states where the enzyme is primed for docking of the E2-Ub complex. To examine which state could accommodate E2 binding, we docked the Nedd4 HECT-Ubch5b crystal structure into both states^25^ (Figure 5A). When docked into state1, the E2 sterically clashes with the RCC1 domain, and although we do observe dynamic movements of the whole dimer, the cleft separating the HECT and RCC1 domains of the protomers does not enlarge sufficiently to accommodate the E2 enzyme. In contrast, the transition of the HECT domain into the state2 creates a wider space compatible with E2 binding. As a subsequent step, the C-lobe rotation towards the N-lobe is necessary for ubiquitin transfer from the E2 to the catalytic cysteine (Figure 5B). This step must be preceded by the plug loop removal, and it remains to be elucidated which conformational change accommodates this requirement.

**Figure 5.**
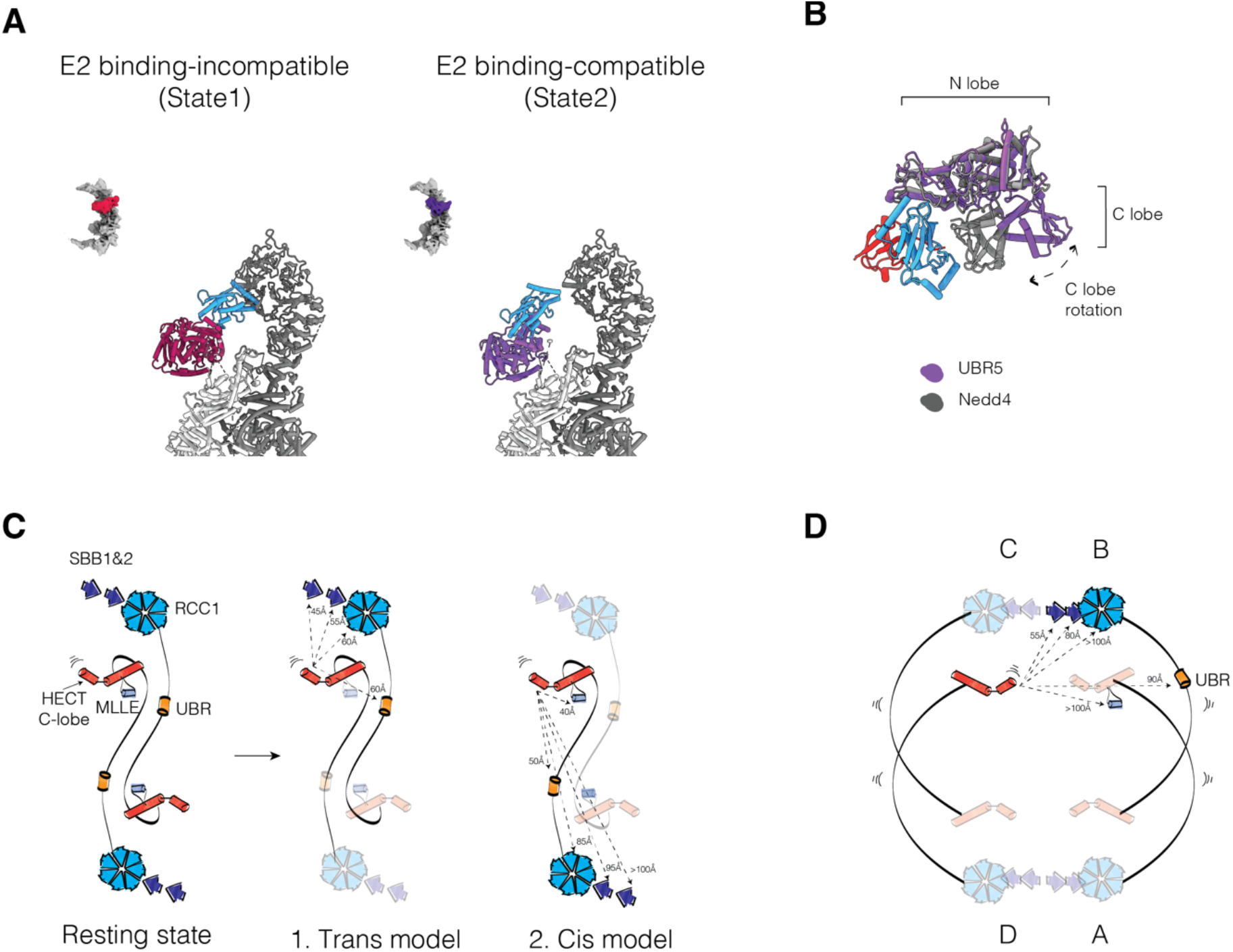
Dynamics of the UBR5 oligomers for E2 enzyme and substrate engagement. A) The two observed states of the HECT domain. The E2 enzyme, Ubch5b (in blue), is docked on the binding site of the HECT domain in either state based on the available crystal structure of Nedd4:Ubch5b:Ubiquitin (PDB:3VJZ). For clarity, only the E2 from this structure is shown. B) An overlay of the UBR5 HECT domain and the crystal structure from A, showing the positions of the C lobe for ubiquitin transfer. The dashed arrow represents the movement of the C lobe for the transfer. Ubiquitin is depicted in red. C) Approximate distances of the protein-interacting motifs to the catalytic cysteine of the UBR5 HECT, in cis and trans. D) Approximate distances of the protein-interacting motifs of one dimer to the catalytic cysteine of a second timer in the tetrameric UBR5 assembly. Protomers of the tetramer are labelled A-D.

E2∼Ub loading and ubiquitin transfer to UBR5’s catalytic cysteine are events not coupled to substrate engagement, but they do not have to exclude each other. Importantly, because of the antiparallel order of the UBR5 protomers, the HECT domains are in closer proximity to the protein-interacting motifs of the second protomer, being separated by about 50Å in the UBR5 resting state (Figure 5C). This suggests that UBR5 ubiquitinates its substrates in a trans fashion: engaging the substrate through one protomer, and mediating ubiquitin transfer with the other. It is also, however, possible that ubiquitination occurs in cis, provided the substrate is large enough to span the distance required. Moreover, the ring architecture of the tetramer positions the catalytic and protein-binding motifs inside the lumen of the ring, opening the possibility that ubiquitination occurs in trans between two opposite dimers (Figure 5D). The architecture and cis/trans ubiquitination mechanisms could further explain the observed adaptability in UBR5’s substrates choices and its broad implications in cellular and cancer biology.

## DISCUSSION

One of the largest E3 ligases, UBR5, is an important regulator of multiple cellular processes with a wide array of substrates. Its exclusive nuclear localisation and implication in cancer biology make it an interesting target for cancer therapies. Our structural and functional characterisation of UBR5 exposed a large helical scaffold decorated with numerous protein-interacting domains, suggesting that there may be multiple modes of substrate recognition with distinct features. Therefore, the potential exists to selectively target UBR5 functions in the different substrate and cellular contexts for cancer therapeutics.

An added complexity of targeting UBR5 is the homodimeric scaffold, a feature that is not commonly reported for HECT-type E3 ligases^41^. Furthermore, we show that the dimers further assemble into tetrameric and higher oligomeric states. While many E3s dimerise, few are reported to form higher oligomeric states, and the function of these higher-order states is unclear. We assigned the oligomerisation properties to the SBB domains, a domain that was previously not reported for UBR5. The elegant assembly of UBR5 protomers into dimers and tetramers allows each interacting domain to lie in close vicinity to a catalytic domain. The stable interactions and conservation scores of the dimeric interface strongly suggest the dimer to be imperative for function, whilst tetramers and higher oligomers are likely to occur. It remains unclear what is the relevance of the tetrameric assembly and whether such assemblies are induced during particular functions or conditions. Speculatively, the large assembly could allow the recruitment of multiple substrates simultaneously or multisubunit complexes. For example, the mitotic checkpoint complex is composed of several components which bind UBR5 directly^4^. Secondly, the large armadillo-type repeat scaffold gives UBR5 conformational flexibility, as is visible in our 3D variability analyses. Both dimeric and tetrameric assemblies showed movements not only of their individual domains but of the whole protein. Such ring-shaped architecture has been observed in several monomeric and multimeric E3 enzymes such as HUWE1, GID or the SCF, with the observations that it promotes substrate recruitment and ubiquitination ^27,28,42,43^.

We identified an interacting partner of UBR5, AKIRIN2. Interestingly, only AKIRIN2 carrying a monoubiquitin modification was efficiently ubiquitinated by UBR5. The chain elongation function of UBR5 is further exhibited when only free ubiquitin was used as the substrate. We further show that the binding of AKIRIN2 and UBR5 is enhanced by its ubiquitin modification. We, therefore, suggest that UBR5 is a chain-elongating E3 ligase which first needs to be presented with a monoubiquitinated substrate or recruited together with other chain-initiating enzymes. The concept of a handoff or sequential mechanism is a recurring theme, especially when branched ubiquitin chains (ubiquitin simultaneously modified at multiple lysines) are formed ^44,45^.

Many questions remain about the polyubiquitination mechanism of UBR5. It is likely that pre-ubiquitinated AKIRIN2 binds one of the protein-interacting motifs of UBR5 in a specific fashion, followed by the engagement of its ubiquitin modification to the UBA domain^23^ (Figure 6). The domain sits on flexible loops and is, therefore, most likely not restricted, sampling a large amount of space in the lumen of the tetrameric ring. Our observations suggest that monoubiquitinated substrates, even with no specific substrate binding site, could engage UBR5 by transient low-affinity binding, making UBR5 a promiscuous, chain-elongating E3 ligase. Supported by the observed similarities to HUWE1 HECT domain flexibility, the dynamic nature of UBR5 assemblies allows the protein-interacting motifs and catalytic domains to be in close proximity between individual protomers. Such arrangements could have enhancing effects on both substrate engagement and chain elongation. While chain elongation is not strictly controlling the fate of a given substrate, it will alter the degradation rates and, consequently the concentration of specific factors. This is consistent with our previous data, which suggests that the absence of UBR5 increases the cellular concentration of MYC but not inducing the apoptosis threshold^1^. These fine-regulating properties make it a likely proto-oncogene as well as a tumour suppressor.

**Figure 6.**
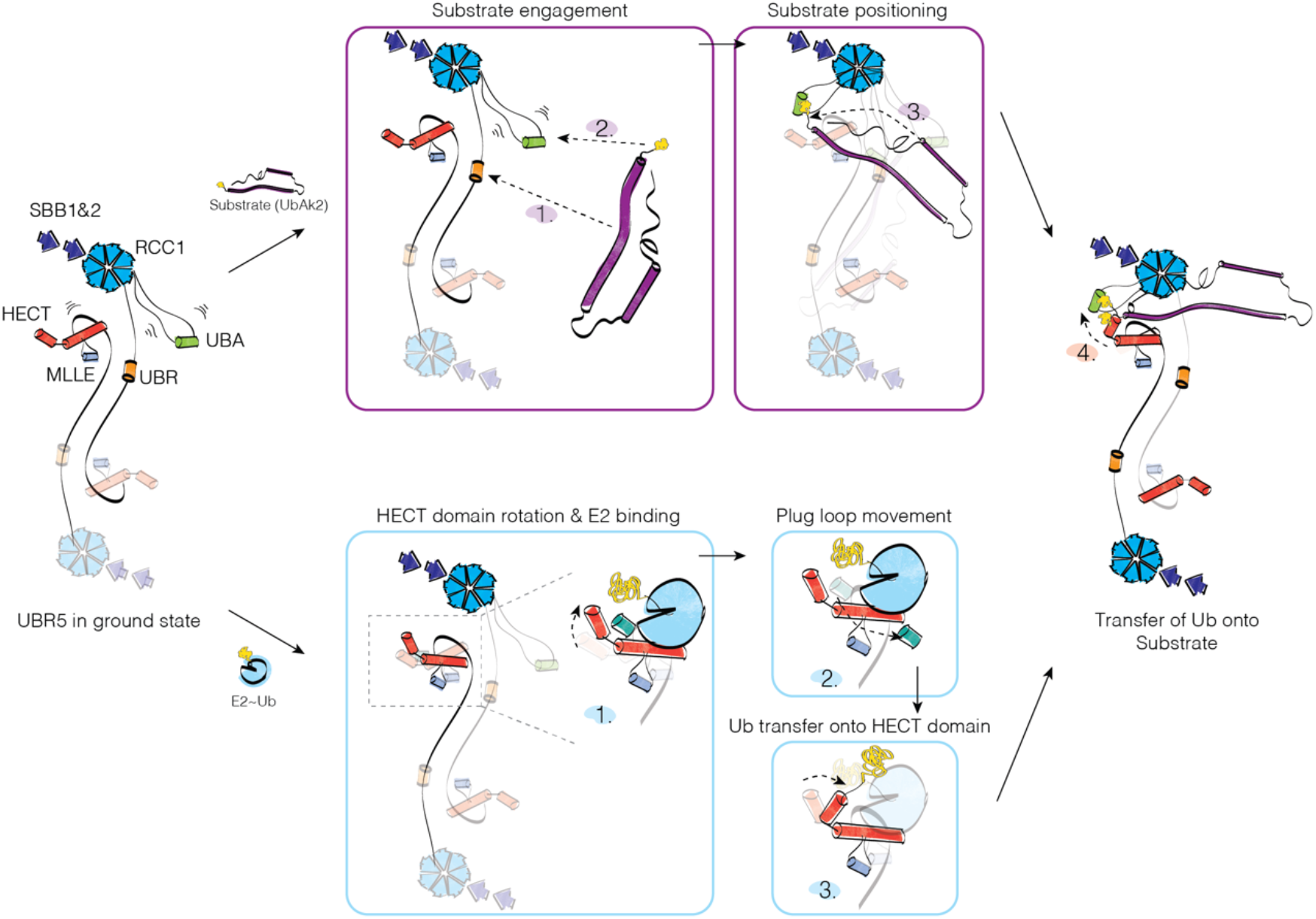
A mechanism of action for UBR5 for substrate engagement and ubiquitin transfer. The substrate is first recruited to one of the protein-interacting motifs of UBR5, followed by its engagement by the UBA domain. Prior to, or in parallel, E2 engagement and ubiquitin transfer from the E2 onto the HECT catalytic cysteine takes place. The acceptor ubiquitin of the substrate is positioned in close proximity to receive the ubiquitin loaded on the HECT domain. Domains are colour-coded as in Figure 1A.

Our findings that AKIRIN2 binds UBR5 but does not get ubiquitinated in its unmodified form raises the question of the reason for their interaction. We previously identified AKIRIN2 as an essential factor for proteasome nuclear import^1^. It is plausible that it performs a similar role for the nuclear localisation of UBR5. Likewise, UBR5 and AKIRIN2 could interact post nuclear import. UBR5, amongst other E3 ligases, has been identified in stress foci together with the proteasome and other factors mediating proteotoxic stress^39,46^. These foci serve as proteolytic centres which cluster UPS components for the rapid degradation of misfolded proteins^47^. It remains to be investigated if UBR5 can interact with the proteasome or if AKIRIN2 could bridge the proteasome with UBR5 in certain conditions.

In conclusion, the presented structural information expands on our knowledge of the activity of HECT E3 ligases, serving as further support for mechanistic studies. The observed similarities and differences to the only other full-length structure, HUWE1, open numerous questions on the regulation of HECT E3 ligase activity by their own architecture. The complex structure and oligomerisation properties of UBR5 begin to explain its broad implications in cancer biology. UBR5 can have inverse effects: in certain cancers, its higher activity could be beneficial for the removal of certain substrates. In other cases, its high activity prevents reaching the apoptotic threshold or triggers immune evasion. Our structure can therefore provide a framework for new therapeutics for UBR5-driven diseases.

## METHODS

### Constructs and cell lines

The codon-optimised sequence of full-length human UBR5 was cloned into pMEPI plasmid as two constructs. For single-particle analysis, the construct included a decahistidine (10xHis) tag followed by a Precission protease (3C) cleavage site, the codon-optimised UBR5 sequence and a double strep tag. For all activity assays, the construct included a double strep tag followed by a 3C site and the codon-optimised UBR5 sequence. The constructs were used for the generation of stable HEK293T cell lines, generated by co-transfecting the constructs under the control of a Tet inducible promoter bearing a puromycin selection marker, and the PiggyBac transposase, for stable genomic integration. Cells were cultured in FreeStyle 293 Expression Medium (GIBCO). Protein expression was induced by the addition of doxycycline at a final concentration of 1 µg/mL for 48 h. 24 h post-induction, 125 mL of Ex-Cell 293 Serum-Free Medium (Sigma Aldrich) was added per 0.5 L of culture. Cells were harvested by centrifugation at 2000x*g* for 20 min. Pellets were washed in 1xPBS and flash-frozen in liquid nitrogen for storage at -70 °C.

The Human AKIRIN2 sequence was cloned into a pGEX vector containing a hexahistidine (6xHis) tag, followed by GFP and a 3C cleavage site. For UbAKIRIN2, the sequence coding for ubiquitin was inserted immediately before the AKIRIN2-coding sequence. For protein expression, BL21(DE3) cells were chemically transformed as per standard protocol. Cells were grown in autoinduction media at 37 °C and expression was induced overnight at 18 °C. Cells were harvested by centrifugation at 2000x*g* for 20 min.

UBCH5B was cloned into a Twist-6xHis-3C plasmid (Twist Bioscience) and expressed in Rosetta2(DE3) competent cells. Cells were grown in autoinduction media at 37 °C, and expression was induced overnight at 18 °C. Cells were harvested by centrifugation at 2000x*g* for 20 min.

Ubiquitin with a deletion of the two C-terminal domains, GG (Ub^ΔGG^), was cloned into a pGEX-TEV vector. For protein expression, BL21(DE3) RIL-competent cells were chemically transformed. Cells were grown in Luria Broth (LB) medium at 37 °C until cells reached OD_600_ 0.8-1.0. Cells were induced with 0.6 mM IPTG and the temperature was reduced to 23 °C overnight.

### Protein purification

For UBR5, the cell pellet was resuspended in 3x (w/v) of BufferA (50 mM HEPES/NaOH pH 8.0, 300 mM NaCl, 0.5 mM TCEP) supplemented with cOmplete protease inhibitor cocktail tablets and Benzonase (IMP Molecular Biology Service) Cells were lysed using a glass Dounce homogeniser and subsequently centrifuged for 1 h at 40,000*g*. The supernatant was applied on a pre-equilibrated StrepTrap HP column (Cytiva) equilibrated in buffer A. The column was washed with 10 column volumes (CV) of BufferA and eluted with BufferA supplemented with 5 mM desthiobiotin. The eluate was collected as fractions and combined, concentrated with a 100 kDa cut-off concentrator (Cytiva), and loaded on a Superose 6 10/300 column (Cytiva) equilibrated in GF Buffer (50mM HEPES/NaOH pH 7.5, 150 mM NaCl, 1 mM TCEP). For long-term storage, the desired fractions were combined, concentrated and flash-frozen in liquid nitrogen. For cryo-EM sample preparation, sample was taken from a single fraction of the SEC elution peak and plunge-frozen directly.

For GFP-AKIRIN2 and GFP-UbAKIRIN2, cell pellet was resuspended in BufferH (50 mM TrisHCl pH 8, 500 mM NaCl, 0.5 mM TCEP) supplemented with cOmplete protease inhibitor cocktail and Benzonase. The cell suspension was homogenised using French Press at 1.5 kBar and subsequently centrifuged for 1 hour at 40,000*g*. The supernatant was applied on a 5 ml HisTrap HP column (Cytiva) pre-equilibrated with buffer H. Column was washed with 10CV of 5 % of buffer E (50 mM TrisHcl pH 8, 500 mM NaCl, 0.5 mM TCEP, 500 mM Imidazole pH 8) and eluted with 50 % of buffer E. Fractions were pooled, concentrated with a 30 kDa cut-off concentrator (Cytiva) and loaded on a Superdex S200 16/60 column (Cytiva) equilibrated in GF Buffer. Desired fractions were pooled, concentrated and flash-frozen in liquid nitrogen for long-term storage.

For UBCH5B, cells were resuspended in BufferU-A (20 mM Tris pH 8.0, 500 mM NaCl, 0.5 mM TCEP) supplemented with cOmplete protease inhibitor cocktail and Benzonase. The cell suspension was homogenised using French Press at 1.5 kBar and subsequently centrifuged for 1 hour at 40,000*g*. The supernatant was applied on a 5 ml HisTrap HP column (Cytiva) pre-equilibrated with BufferU-A, and protein eluted using a gradient of Buffer U-A and BufferU-B (BufferA supplemented with 500mM Imidazole pH 8). The tag was cleaved overnight using Precission protease and subsequently passed through HisTrap HP column to capture the cleaved tag and protease. Protein was concentrated and loaded onto a Superdex 75 16/60 column (Cytiva) pre-equilibrated with BufferU-GF (20 mM HEPES pH 8.0, 50 mM NaCl, 1 mM DTT). Desired fractions were pooled, concentrated and flash-frozen in liquid nitrogen for long-term storage.

For wild-type ubiquitin (wtUbiquitin), cells were resuspended in BufferUbi (50 mM Tris 7.5, 300 mM NaCl) supplemented with cOmplete protease inhibitor cocktail and Benzonase. The cell suspension was homogenised using French Press at 1.5 kBar and subsequently centrifuged for 1 hour at 40,000*g*. Glacial acetic acid was slowly added to the supernatant until reaching pH 4.5, followed by centrifugation at 15,000x*g* for 30 minutes. The supernatant was then dialysed overnight into 25mM sodium acetate (NaAc) pH 4.5. Dialysed sample was concentrated and loaded on a pre-equilibrated Resource S column (Cytiva) in 25mM NaAc pH 4.5. Protein was eluted with a shallow gradient of 25mM NaAc pH 4.5, and 25mM NaAc pH4.5 and 250mM NaCl. Desired fractions were concentrated using a 5kDa cut-off concentrator (Cytiva) and loaded on a Superdex 75 16/60 column (Cytiva) pre-equilibrated with buffer composed of 25mM HEPES pH 8 and 50mM NaCl.

For Ub^ΔGG^, cells were pelleted at 3000xg, resuspended in BufferT (50 mM Tris pH 7.6, 200mM NaCl), and lysed by sonication. The lysate was cleared by centrifugation at 15000x*g* and incubated with Anti-GS4B resin for 2 h at 4 °C (Genessee Scientific). The resin was washed with excess BufferT and TEV protease was added to beads rocking overnight at RT. The resin was pelleted at 500x*g*, supernatant containing Ub^ΔGG^ was removed and stored, and resin was resuspended in a small amount of BufferT. Excess Ub^ΔGG^ was washed off of resin with more BufferT. Supernatant and wash were combined, centrifuged at 4000x*g* to remove precipitated protein, concentrated, and purified using Ni-NTA Superflow beads (Qiagen) and GS4B resins to final purity. Protein was concentrated and flash-frozen in liquid nitrogen for long-term storage.

Labelling of Ub^ΔGG^ was performed as described previously^48^. Briefly, the protein was reduced by adding 20 mM DTT and desalted twice into 50 mM HEPES pH 7.0, 200 mM NaCl. Cy5-maleimide was resuspended in DMSO and added in 2.5 molar excess to the protein, followed by 2 h incubation at RT. Cy5-Ub^ΔGG^ was dialysed overnight against 50 mM HEPES pH 8.0, 200 mM NaCl, followed by desalting to remove excess dye. The labelled protein was stored at -80 °C.

Securin purification and fluorescent labelling was performed as described previously^49^. Recombinant human UBA1 was provided by the Vienna BioCenter core facilities and purified as described elsewhere ^28^.

### Ubiquitination assays

All reactions were prepared by mixing reaction components in assay buffer (100 mM NaCl, 20 mM HEPES pH 8, 5 mM MgCl_2_) on ice. Reactions were performed at 37 °C with the following final concentrations: HisUBA1 at 0.3 µM, UBCH5b at 2.5 µM, UBR5 at 0.3 µM, wtUbiquitin at 20 µM, the substrate at 2 µM. Reactions were initiated by the addition of 2 mM ATP and collected at the indicated time points. Timepoint zero was taken prior to addition of ATP. Substrate ubiquitination was visualised using fluorescence imaging using a ChemiDoc MP System (BioRad). Gels were subsequently stained with Coomassie blue to visualise protein content.

E2 screen (E2 Scan Kit version 2, Ubiquigent) was performed guided by the manufacturer’s instructions. Briefly, concentrations of components of the reactions were used as described above, except that a mixture of wtUbiquitin and ^FAM^ubiquitin was used to visualise autoubiquitination. Incubation was performed at 37°C for 30minutes. Results were visualized using a ChemiDocMP System (BioRad) using fluorescence imaging and stain-free imaging.

Density gradient centrifugations were performed by preparing the low-density solution of 10 % sucrose and high-density solution of 30 % sucrose in GF buffer. Gradients were made by mixing the two solutions using the Gradient Master (Biocomp systems) to create a continuous sucrose gradient. The sample was applied onto the density gradient prepared in an open-top 4.2ml ultracentrifuge tube and run for 16 h at 28,000 rpm in an SW60Ti rotor. All proteins were loaded at a concentration of 40 pmol. Gradients were manually fractionated into 200 µl fractions and analysed with stain-free and fluorescence imaging using ChemiDocMP System (BioRad)

### Protein sequence analysis

Sequences were collected from the UniProt reference database ^50^in a NCBI-Blast search applying highly significant E-value thresholds (<1e-50)^51^ and aligned with MAFFT (v7.505, -linsi method)^52^. Visualization was performed with Jalview ^53^.

### Cryo-EM sample preparation and data collection

Open-hole R1.2/1.3 grids (Quantifoil) with 200 mesh were used for plunge freezing. Grids were glow-discharged for 120 s at 25 mA using residual air. 4 µl of the sample was applied onto the treated side and front-side blotted using the LeicaGP plunge freezer. UBR5 was frozen in GF Buffer with the addition of 4 mM CHAPSO or 0.005 % fluorinated octyl β-maltoside (Anatrace) prior to freezing.

Initial grid screening was performed on the 200kV Glacios microscope (ThermoFisher) using the Falcon III detector (ThermoFisher). For high-resolution structure determination, data collection was performed on a 300 kV Titan Krios G4 equipped with a cold field emission gun and a post-column Selectris energy filter (ThermoFisher) with a 5 eV slit width and a Falcon 4 or Falcon 4i direct electron detector (ThermoFisher). Images were collected at a pixel size of 0.745 Å/pix or 0.951Å/pix with a cumulative dose of 40e^−^/Å^2^ for untilted, and 50e-/Å2 for 30° tilted images, in eer format, using a defocus range in 0.3µm increments (Table1).

### Cryo-EM image processing and model refinement

On-the-fly preprocessing (patch motion correction and CTF estimation) was performed using Cryosparc Live. Particles were manually picked in Cryosparc 3.1^54^ to generate templates for template picking, which was performed on manually curated micrographs. Following template-based picking, particles were classified using reference-free 2D classification, followed by Ab-Initio reconstruction and heterogeneous refinement. The best resulting 3D volume was subjected to several rounds of homogeneous and local refinements using half-map masks generated in ChimeraX. Final models were sharpened using the DeepEMhancer tool^55^. The initial model for model building was generated using the AlphaFold2 prediction software ^56^. The predicted model was docked twice onto the maps and manually edited using Coot and ISOLDE^57^. Real-space refinement was performed using Phenix^58^.

### Mass Photometry

Measurements were performed as described in^29^. Briefly, OneMP (Refeyn) was calibrated using the Native protein marker as a protein standard using a medium field of view. MP signals were collected for 60 s to detect at least 5000 individual molecules. Raw data was processed using the DiscoverMP software and plotted as histograms of molar mass distributions.

## ACKNOWLEDGEMENTS

We thank all members of the Haselbach and Brown lab for discussions on the project, the Electron Microscopy, ProTech and Mass Spectrometry facilities from the Vienna Biocenter Core Facilities for their support. Z.H. received funding from the European Union’s Framework Programme for Research and Innovation Horizon 2020 (2014-2020) under the Marie Curie Skłodowska Grant Agreement Nr. 847548. H.L.B. was generously supported by Boehringer Ingelheim (BI1982). IMP is supported by Boehringer Ingelheim. The Brown lab is supported by NIH T32GM008570 (D.L.B) and NIH R35GM128855 (N.G.B.).

## AUTHOR CONTRIBUTIONS

D.H, N.G.B and Z.H. conceived and planned the project. I.G. purified UBR5 proteins and prepared cryo-EM grids. I.G and Z.H performed grid screening and biochemical assays. H.K collected datasets for high-resolution structure determination. Z.H performed cryo-EM analysis. Z.H and D.H built the molecular model. I.G, H.L.B and D.L.B. purified proteins for assays. K.B assisted with OneMP measurements. A.S performed bioinformatic analysis. Z.H, N.G.B and D.H prepared the manuscript, with input from all authors.

## DATA AVAILABILITY

Atomic coordinates and cryo-EM density maps have been deposited in the Protein Data Bank (PDBe) under accession codes PDB:8BJA/EMD:16087. The raw micrographs were submitted to the EMPIAR database (EMPIAR-XXXXX). Other source data are included in the paper.

## SUPPLEMENTAL INFORMATION

**Supplemental Figure 1.**
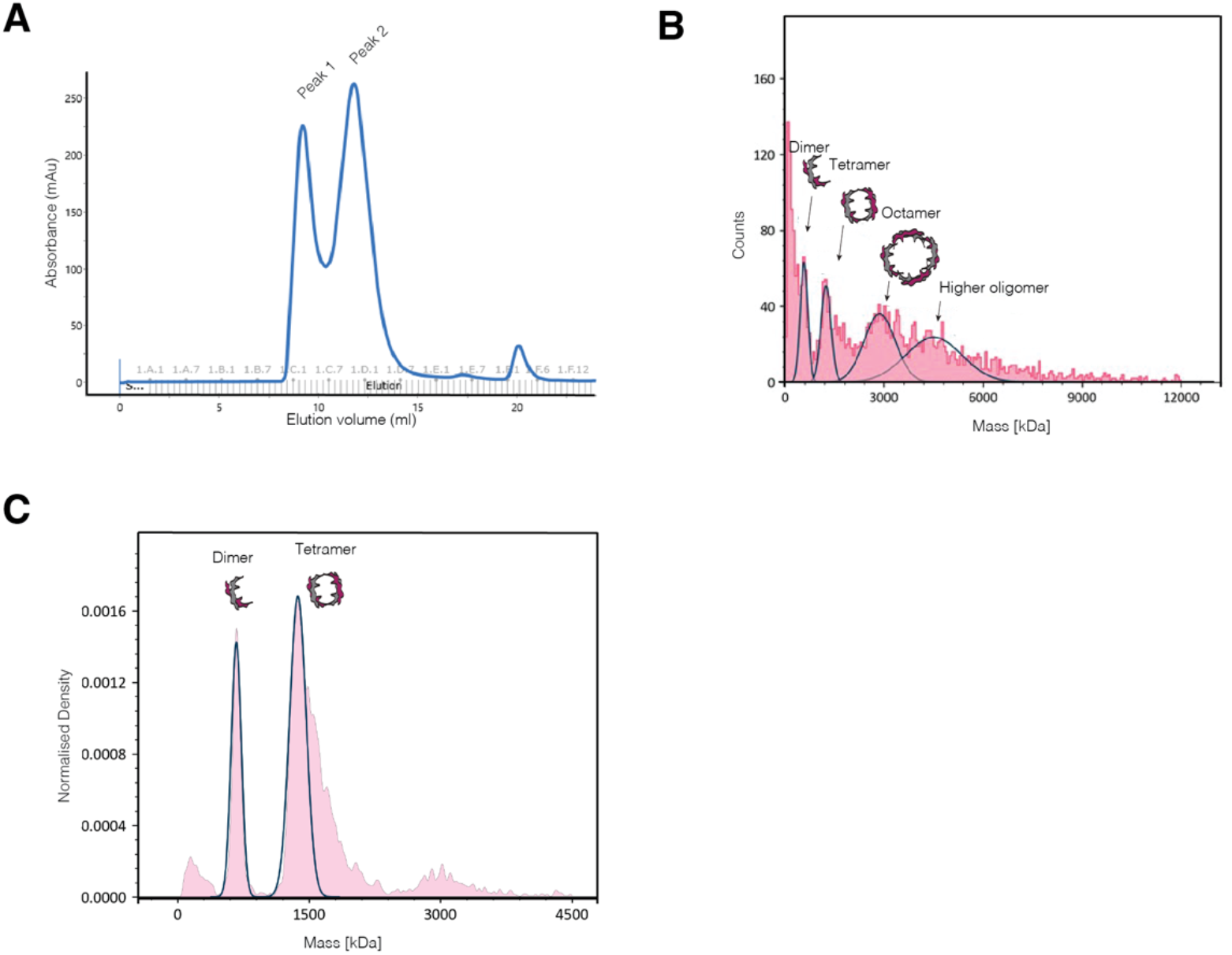
Purification analysis of the recombinant UBR5 protein. A) Size exclusion chromatography elution profile. B) Mass distribution and oligomeric states adopted by UBR5 in the first elution peak from A, measured using mass photometry. C) Mass distribution of UBR5 diluted in a high concentration of NaCl, measured using mass photometry.

**Supplemental Figure 2.**
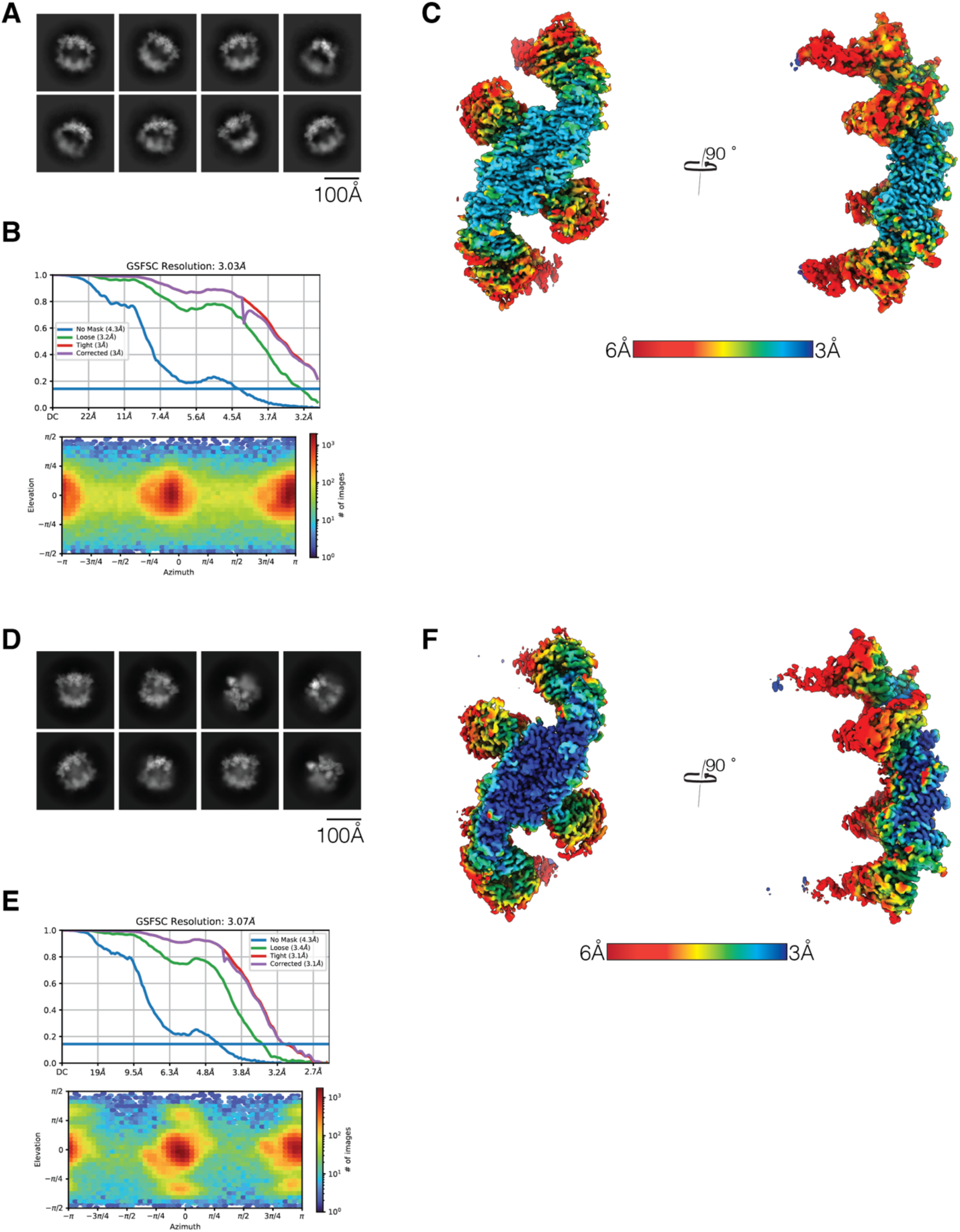
Resolution and angle distributions of the UBR5 Cryo-EM maps. A) Representative 2D class averages of Map 1 (EMDB--). B) Final resolution of Map 1 as estimated by the FSC curve and angular distribution of the final map. C) Local resolution of Map 1. D) Representative 2D class averages of Map 2. E) Final resolution of Map 2 as estimated by the FSC curve and angular distribution of the final map. F) Local resolution of Map 2.

**Supplemental Figure 3.**
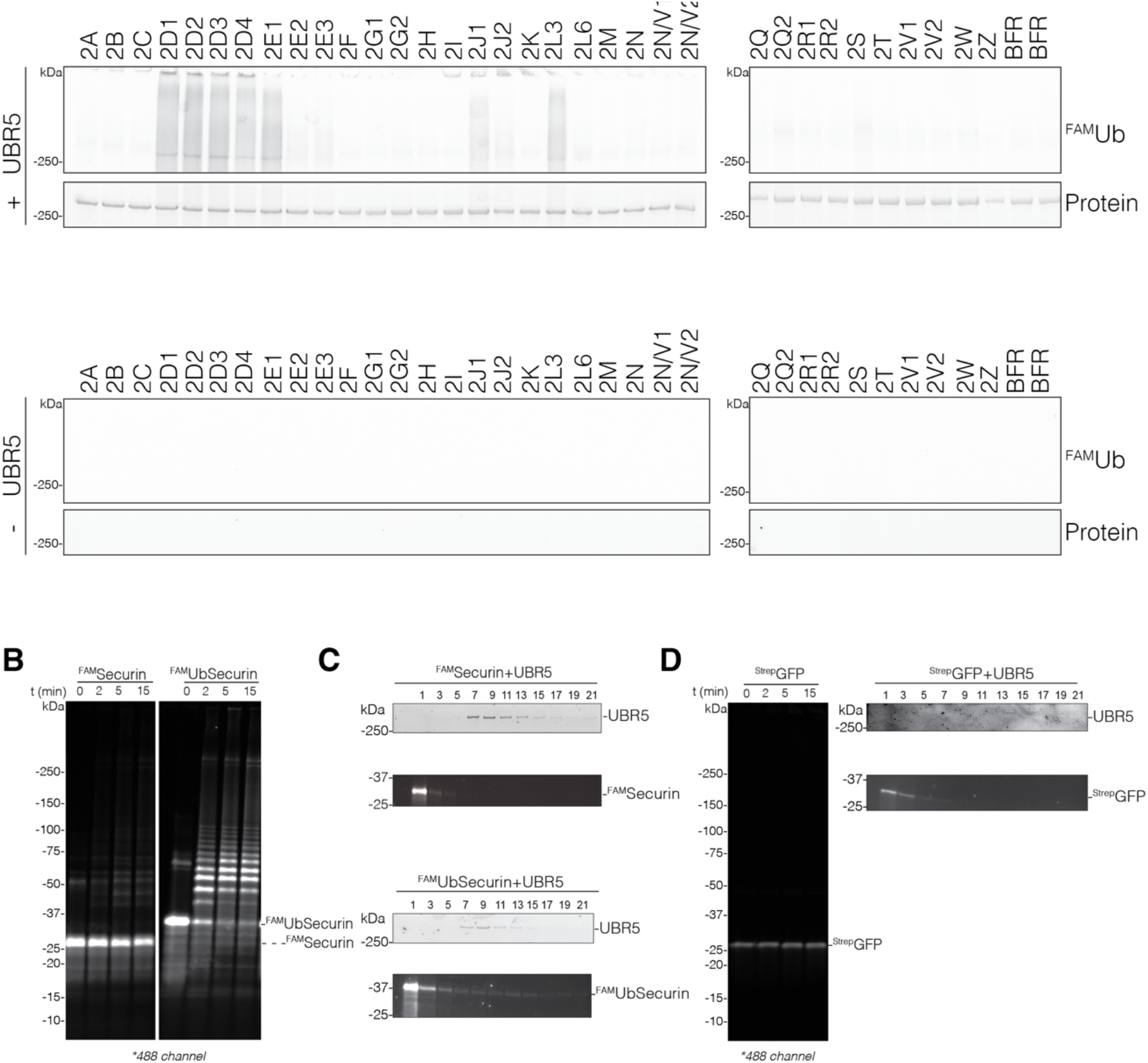
E2 selection and substrate ubiquitination and selection of UBR5. (Related to Figure 3) A) E2 screening kit. Top panel contains conditions supplemented with UBR5. Fluorescence imaging was used to visualise UBR5 auto-ubiquitination, and total protein levels visualised by stain-free imaging. Bottom panel shows conditions unsupplemented with UBR5, imaged using the same fluorescence and stain-free imaging parameters. B) Ubiquitination assays of an unmodified and monoubiquitin-like nonspecific substrate. Ubiquitinated products were visualised by imaging the FAM label. C) Sucrose gradient sedimentation profiles of UBR5 mixed with two control substrates. UBR5 was visualised using stain-free imaging. Substrates were visualised by imaging their FAM label. D) Sucrose gradients and ubiquitination of the GFP tag of AKIRIN2. Same conditions as in C were used for imaging.

**Supplemental Table1.**
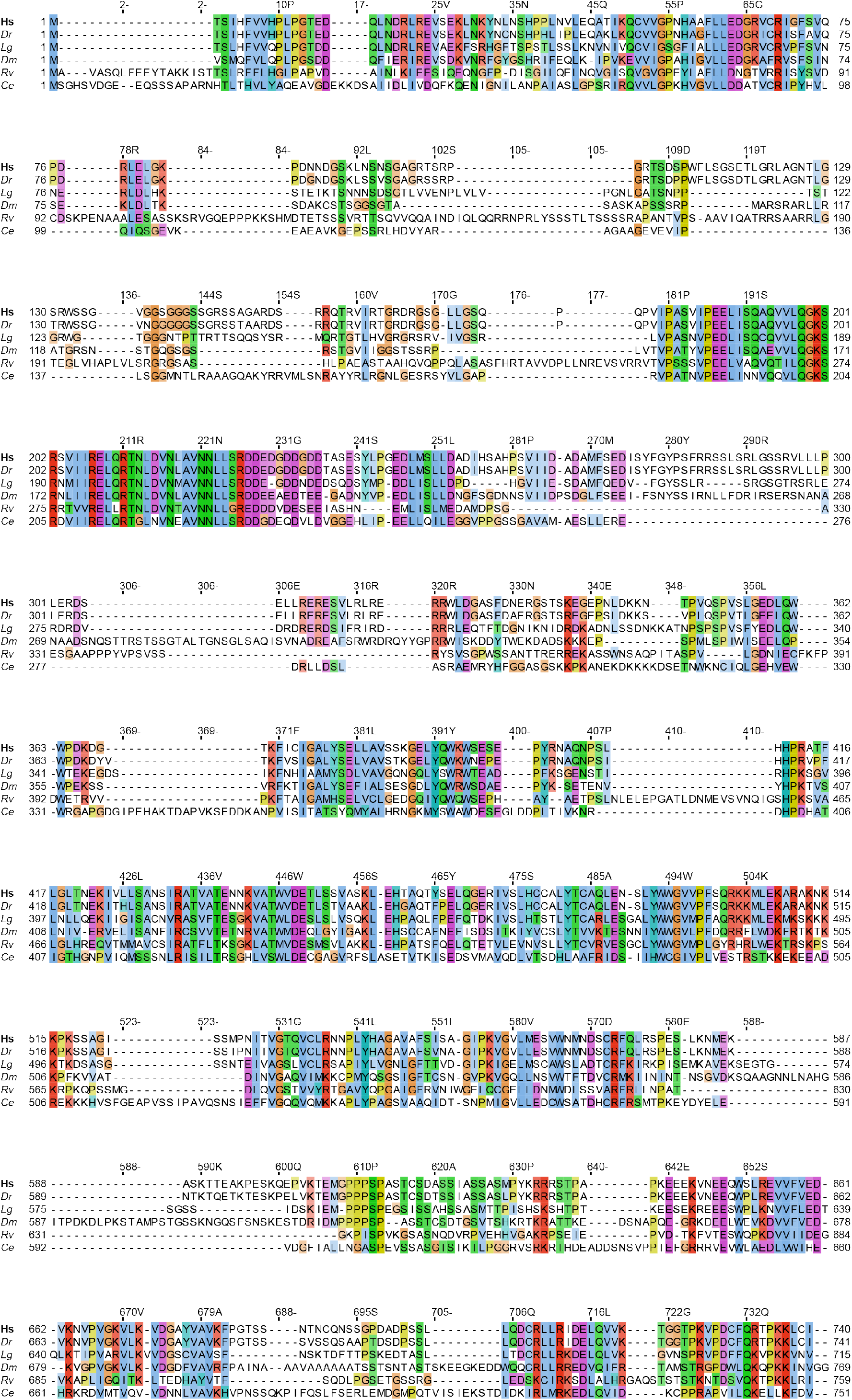
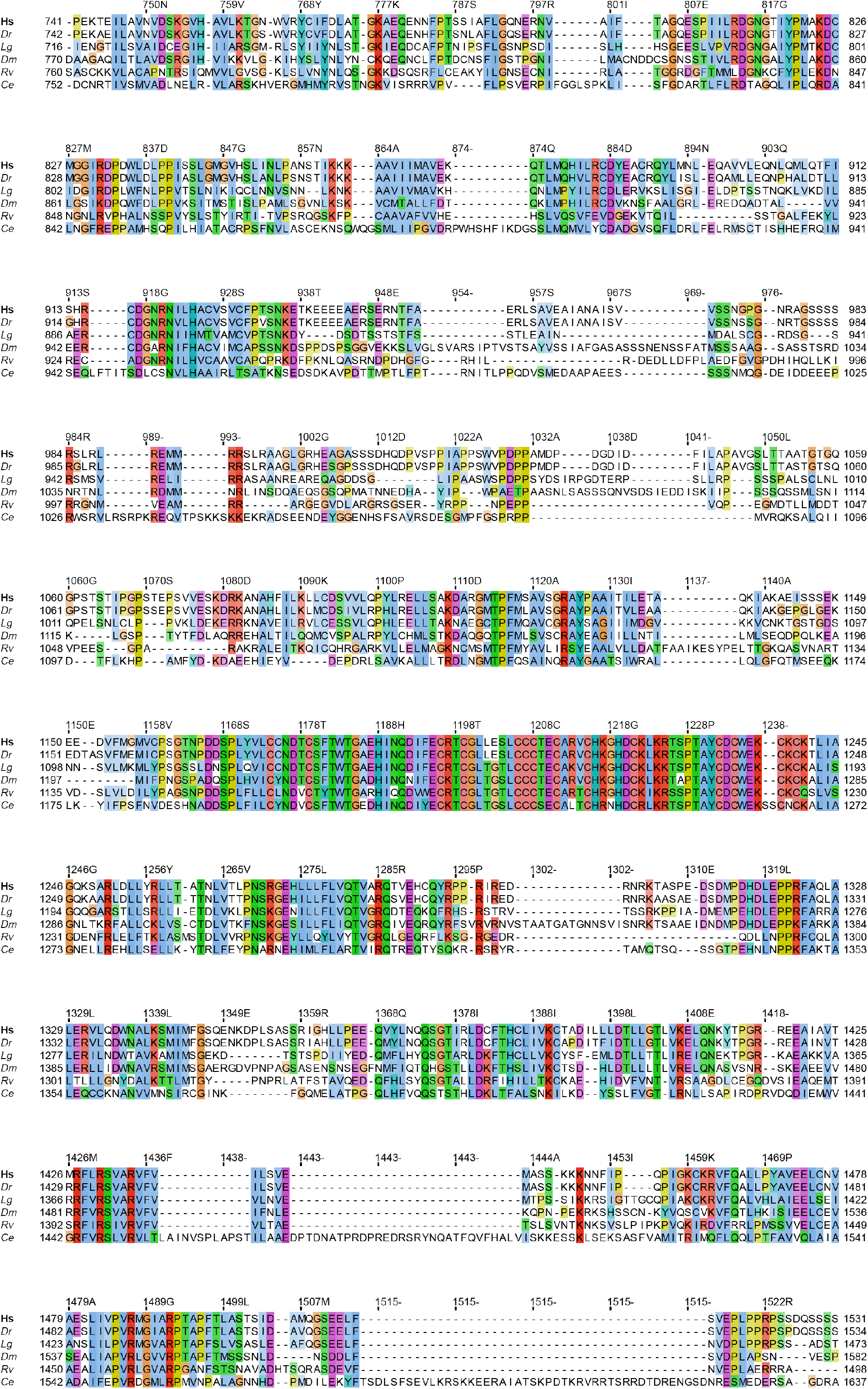
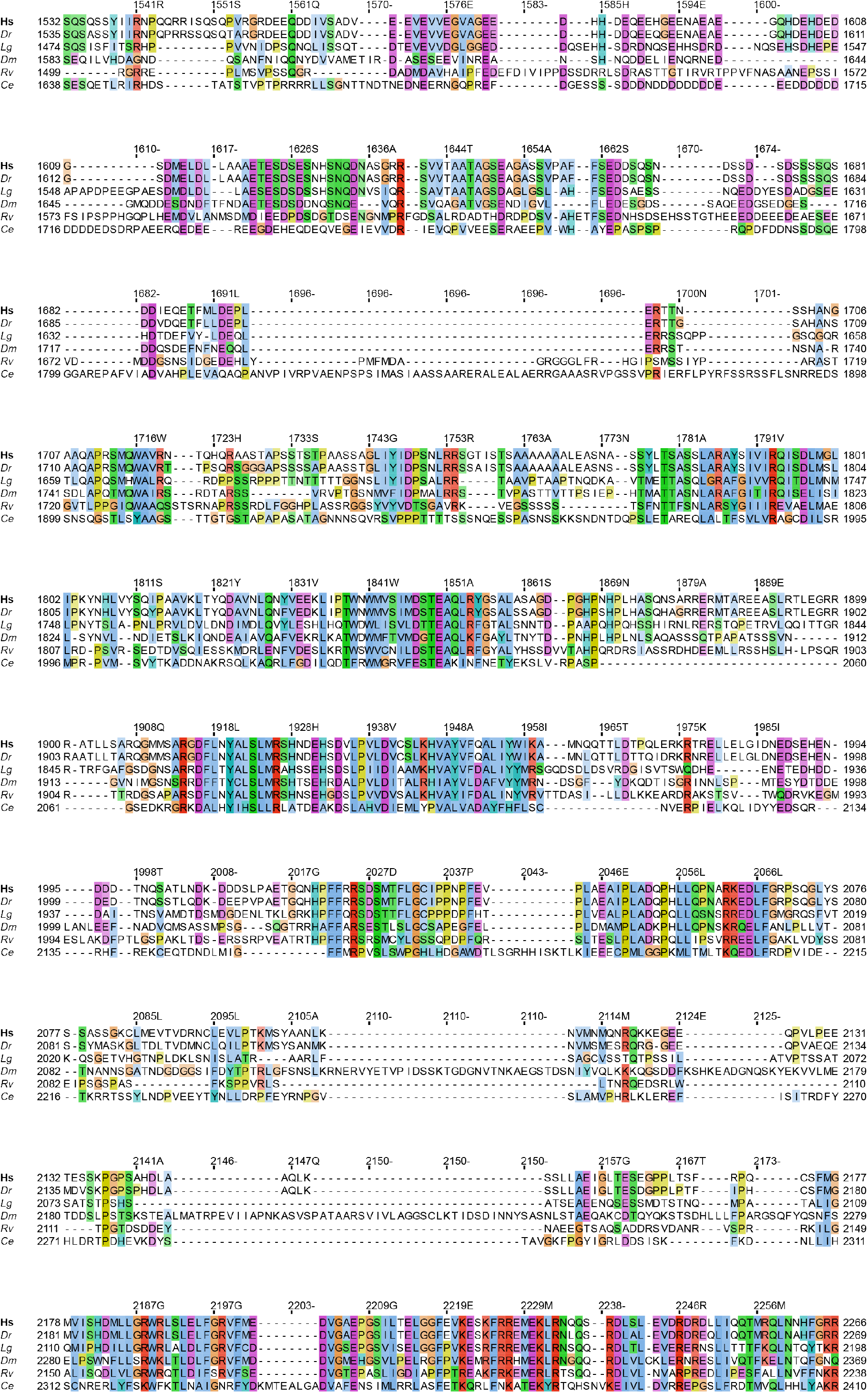
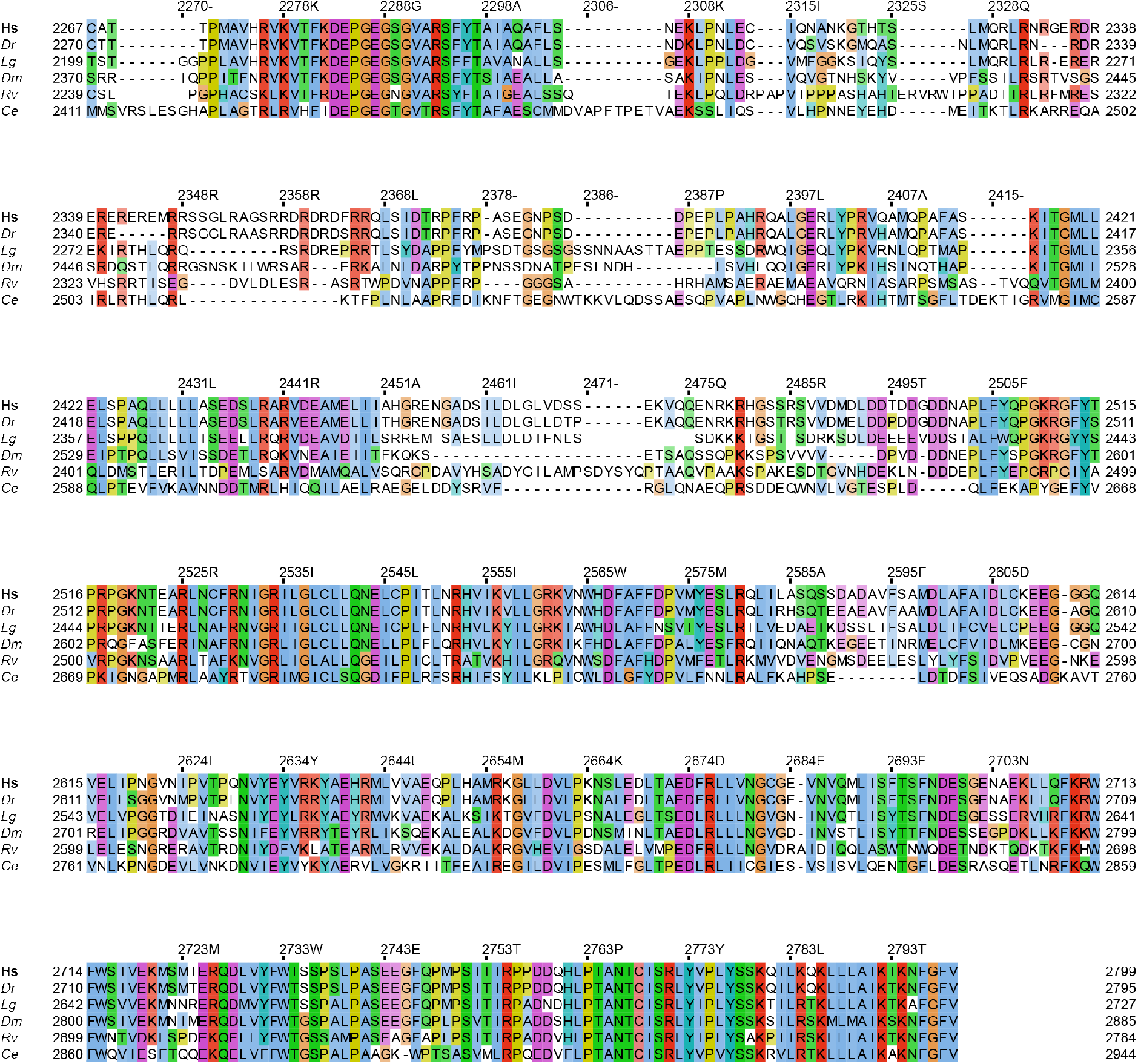
Sequence alignments and conservation of UBR5 across species. Uniprot IDs are shown in brackets. *Hs - Homo sapiens [Q95701], Dr - Danio rerio [F1Q8J2], Lg - Lottia gigantea [V3Z5H7], Dm - Drosophila melanogaster [P51592], Rv - Ramazzottius varieornatus [A0A1D1V0S6], Ce - Caenorhabditis elegans [G5EDT9]*.

**Supplemental Table2. Ubiquitin chains formed by UBR5.** Mass spectrometry data of ubiquitin chain types identified in ubiquitination assays by UBR5. Data was obtained from experiment shown in Fig.3C.

## REFERENCES

1. de Almeida, M. et al. AKIRIN2 controls the nuclear import of proteasomes in vertebrates. Nature 599, 491–496 (2021).

2. Mansfield, E., Hersperger, E., Biggs, J. & Shearn, A. Genetic and Molecular Analysis of hyperplastic discs, a Gene Whose Product Is Required for Regulation of Cell Proliferation in Drosophila melanogaster Imaginal Discs and Germ Cells. Dev. Biol. 165, 507–526 (1994).

3. Scialpi, F., Mellis, D. & Ditzel, M. EDD, a ubiquitin-protein ligase of the N-end rule pathway, associates with spindle assembly checkpoint components and regulates the mitotic response to nocodazole. J. Biol. Chem. 290, 12585–12594 (2015).

4. Kaisari, S. et al. Role of ubiquitin-protein ligase UBR5 in the disassembly of mitotic checkpoint complexes. Proc. Natl. Acad. Sci. U. S. A. 119, (2022).

5. de Vivo, A. et al. The OTUD5-UBR5 complex regulates FACT-mediated transcription at damaged chromatin. Nucleic Acids Res. 47, 729–746 (2019).

6. Henderson, M. J. et al. EDD mediates DNA damage-induced activation of CHK2. J. Biol. Chem. 281, 39990–40000 (2006).

7. Zhang, T., Cronshaw, J., Kanu, N., Snijders, A. P. & Behrens, A. UBR5-mediated ubiquitination of ATMIN is required for ionizing radiation-induced ATM signaling and function. Proc. Natl. Acad. Sci. U. S. A. 111, 12091–12096 (2014).

8. Gudjonsson, T. et al. TRIP12 and UBR5 suppress spreading of chromatin ubiquitylation at damaged chromosomes. Cell 150, 697–709 (2012).

9. Cammarata-Mouchtouris, A. et al. Hyd ubiquitinates the NF-κB co-factor Akirin to operate an effective immune response in Drosophila. PLoS Pathog. 16, e1008458 (2020).

10. Shearer, R. F., Iconomou, M., Watts, C. K. W. & Saunders, D. N. Functional Roles of the E3 Ubiquitin Ligase UBR5 in Cancer. Mol. Cancer Res. 13, 1523–1532 (2015).

11. Ling, S. & Lin, W.-C. EDD inhibits ATM-mediated phosphorylation of p53. J. Biol. Chem. 286, 14972–14982 (2011).

12. Flack, J. E., Mieszczanek, J., Novcic, N. & Bienz, M. Wnt-Dependent Inactivation of the Groucho/TLE Co-repressor by the HECT E3 Ubiquitin Ligase Hyd/UBR5. Mol. Cell 67, 181-193.e5 (2017).

13. Qiao, X. et al. UBR5 Is Coamplified with MYC in Breast Tumors and Encodes an Ubiquitin Ligase That Limits MYC-Dependent Apoptosis. Cancer Res. 80, 1414– 1427 (2020).

14. Schukur, L. et al. Identification of the HECT E3 ligase UBR5 as a regulator of MYC degradation using a CRISPR/Cas9 screen. Sci. Rep. 10, 20044 (2020).

15. Liao, L. et al. E3 Ubiquitin Ligase UBR5 Drives the Growth and Metastasis of Triple-Negative Breast Cancer. Cancer Res. 77, 2090–2101 (2017).

16. Clancy, J. L. et al. EDD, the human orthologue of the hyperplastic discs tumour suppressor gene, is amplified and overexpressed in cancer. Oncogene 22, 5070– 5081 (2003).

17. Li, J. et al. E3 Ubiquitin Ligase UBR5 Promotes the Metastasis of Pancreatic Cancer via Destabilizing F-Actin Capping Protein CAPZA1. Front. Oncol. 11, 634167 (2021).

18. Wang, J., Zhao, X., Jin, L., Wu, G. & Yang, Y. UBR5 Contributes to Colorectal Cancer Progression by Destabilizing the Tumor Suppressor ECRG4. Dig. Dis. Sci. 62, 2781–2789 (2017).

19. Meissner, B. et al. The E3 ubiquitin ligase UBR5 is recurrently mutated in mantle cell lymphoma. Blood 121, 3161–3164 (2013).

20. Song, M. et al. Tumor derived UBR5 promotes ovarian cancer growth and metastasis through inducing immunosuppressive macrophages. Nat. Commun. 11, 6298 (2020).

21. Song, M. et al. Targeting ubiquitin protein ligase E3 component N-recognin 5 in cancer cells induces a CD8+ T cell mediated immune response. Oncoimmunology 9, 1746148 (2020).

22. Muñoz-Escobar, J., Matta-Camacho, E., Kozlov, G. & Gehring, K. The MLLE domain of the ubiquitin ligase UBR5 binds to its catalytic domain to regulate substrate binding. J. Biol. Chem. 290, 22841–22850 (2015).

23. Kozlov, G. et al. Structural basis of ubiquitin recognition by the ubiquitin-associated (UBA) domain of the ubiquitin ligase EDD. J. Biol. Chem. 282, 35787– 35795 (2007).

24. Lorenz, S. Structural mechanisms of HECT-type ubiquitin ligases. Biol. Chem. 399, 127–145 (2018).

25. Kamadurai, H. B. et al. Insights into ubiquitin transfer cascades from a structure of a UbcH5B approximately ubiquitin-HECT(NEDD4L) complex. Mol. Cell 36, 1095–1102 (2009).

26. Maspero, E. et al. Structure of the HECT:ubiquitin complex and its role in ubiquitin chain elongation. EMBO Rep. 12, 342–349 (2011).

27. Hunkeler, M. et al. Solenoid architecture of HUWE1 contributes to ligase activity and substrate recognition. Mol. Cell 81, 3468-3480.e7 (2021).

28. Grabarczyk, D. B. et al. HUWE1 employs a giant substrate-binding ring to feed and regulate its HECT E3 domain. Nat. Chem. Biol. 17, 1084–1092 (2021).

29. Sonn-Segev, A. et al. Quantifying the heterogeneity of macromolecular machines by mass photometry. Nat. Commun. 11, 1772 (2020).

30. Renault, L., Kuhlmann, J., Henkel, A. & Wittinghofer, A. Structural basis for guanine nucleotide exchange on Ran by the regulator of chromosome condensation (RCC1). Cell 105, 245–255 (2001).

31. Youkharibache, P. et al. The Small β-Barrel Domain: A Survey-Based Structural Analysis. Structure 27, 6–26 (2019).

32. Punjani, A. & Fleet, D. J. 3D variability analysis: Resolving continuous flexibility and discrete heterogeneity from single particle cryo-EM. J. Struct. Biol. 213, 107702 (2021).

33. Varshavsky, A. The N-end rule pathway and regulation by proteolysis. Protein Sci. 20, 1298–1345 (2011).

34. Kim, J. G. et al. Signaling Pathways Regulated by UBR Box-Containing E3 Ligases. Int. J. Mol. Sci. 22, (2021).

35. Choi, W. S. et al. Structural basis for the recognition of N-end rule substrates by the UBR box of ubiquitin ligases. Nat. Struct. Mol. Biol. 17, 1175–1181 (2010).

36. Matta-Camacho, E., Kozlov, G., Li, F. F. & Gehring, K. Structural basis of substrate recognition and specificity in the N-end rule pathway. Nat. Struct. Mol. Biol. 17, 1182–1187 (2010).

37. Muñoz-Escobar, J., Matta-Camacho, E., Cho, C., Kozlov, G. & Gehring, K. Bound Waters Mediate Binding of Diverse Substrates to a Ubiquitin Ligase. Structure 25, 719-729.e3 (2017).

38. Tasaki, T. et al. The substrate recognition domains of the N-end rule pathway. J. Biol. Chem. 284, 1884–1895 (2009).

39. Yau, R. G. et al. Assembly and Function of Heterotypic Ubiquitin Chains in Cell-Cycle and Protein Quality Control. Cell 171, 918-933.e20 (2017).

40. Ohtake, F., Tsuchiya, H., Saeki, Y. & Tanaka, K. K63 ubiquitylation triggers proteasomal degradation by seeding branched ubiquitin chains. Proc. Natl. Acad. Sci. U. S. A. 115, E1401–E1408 (2018).

41. Ronchi, V. P., Klein, J. M., Edwards, D. J. & Haas, A. L. The active form of E6-associated protein (E6AP)/UBE3A ubiquitin ligase is an oligomer. J. Biol. Chem. 289, 1033–1048 (2014).

42. Sherpa, D. et al. GID E3 ligase supramolecular chelate assembly configures multipronged ubiquitin targeting of an oligomeric metabolic enzyme. Mol. Cell 81, 2445-2459.e13 (2021).

43. Welcker, M. et al. Two diphosphorylated degrons control c-Myc degradation by the Fbw7 tumor suppressor. Sci Adv 8, eabl7872 (2022).

44. Brown, N. G. et al. Dual RING E3 Architectures Regulate Multiubiquitination and Ubiquitin Chain Elongation by APC/C. Cell 165, 1440–1453 (2016).

45. Haakonsen, D. L. & Rape, M. Branching Out: Improved Signaling by Heterotypic Ubiquitin Chains. Trends Cell Biol. 29, 704–716 (2019).

46. Besche, H. C., Haas, W., Gygi, S. P. & Goldberg, A. L. Isolation of mammalian 26S proteasomes and p97/VCP complexes using the ubiquitin-like domain from HHR23B reveals novel proteasome-associated proteins. Biochemistry 48, 2538– 2549 (2009).

47. Yasuda, S. et al. Stress- and ubiquitylation-dependent phase separation of the proteasome. Nature 578, 296–300 (2020).

48. Brown, N. G. et al. Mechanism of polyubiquitination by human anaphase-promoting complex: RING repurposing for ubiquitin chain assembly. Mol. Cell 56, 246–260 (2014).

49. Welsh, K. A. et al. Functional conservation and divergence of the helix-turn-helix motif of E2 ubiquitin-conjugating enzymes. EMBO J. 41, e108823 (2022).

50. UniProt Consortium. UniProt: the universal protein knowledgebase in 2021. Nucleic Acids Res. 49, D480–D489 (2021).

51. Altschul, S. F. et al. Gapped BLAST and PSI-BLAST: a new generation of protein database search programs. Nucleic Acids Res. 25, 3389–3402 (1997).

52. Katoh, K. & Toh, H. Recent developments in the MAFFT multiple sequence alignment program. Brief. Bioinform. 9, 286–298 (2008).

53. Waterhouse, A. M., Procter, J. B., Martin, D. M. A., Clamp, M. & Barton, G. J. Jalview Version 2--a multiple sequence alignment editor and analysis workbench. Bioinformatics 25, 1189–1191 (2009).

54. Punjani, A., Rubinstein, J. L., Fleet, D. J. & Brubaker, M. A. cryoSPARC: algorithms for rapid unsupervised cryo-EM structure determination. Nat. Methods 14, 290–296 (2017).

55. Sanchez-Garcia, R. et al. DeepEMhancer: a deep learning solution for cryo-EM volume post-processing. Communications Biology 4, 1–8 (2021).

56. Jumper, J. et al. Highly accurate protein structure prediction with AlphaFold. Nature 596, 583–589 (2021).

57. Croll, T. I. ISOLDE: a physically realistic environment for model building into low-resolution electron-density maps. Acta Crystallogr D Struct Biol 74, 519–530 (2018).

58. Afonine, P. V. et al. Real-space refinement in PHENIX for cryo-EM and crystallography. Acta Crystallogr D Struct Biol 74, 531–544 (2018).

